# Phylogeography of bar-tailed godwits: pre-LGM structure in Beringia and westward colonization of post-glacial Europe

**DOI:** 10.1101/2024.04.22.590527

**Authors:** Jesse R. Conklin, Yvonne I. Verkuil, Margaux J.M. Lefebvre, Phil F. Battley, Roeland A. Bom, Robert E. Gill, Chris J. Hassell, Job ten Horn, Daniel R. Ruthrauff, T. Lee Tibbitts, Pavel S. Tomkovich, Nils Warnock, Theunis Piersma, Michaël C. Fontaine

## Abstract

In migratory birds, high mobility may reduce population structure through increased dispersal and enable adaptive responses to environmental change, whereas rigid migratory routines predict low dispersal, increased geographic structure, and limited flexibility to respond to change. We used nextRAD sequencing of 14,318 single-nucleotide polymorphisms to explore the population genetics and phylogeographic history of the bar-tailed godwit, *Limosa lapponica*, a migratory shorebird with six recognized subspecies and known for making the longest non-stop flights of any landbird. Using scenario-testing in an Approximate Bayesian Computation framework, we infer that bar-tailed godwits existed in three main lineages at the Last Glacial Maximum (LGM), when much of their present-day Arctic and sub-Arctic breeding range persisted in a large, unglaciated Siberian-Beringian refugium. Subsequently, population structure developed at both longitudinal extremes: in the east, a genetic cline exists across latitude in the Alaska breeding range of subspecies *L. l. baueri*; in the west, one lineage diversified into three extant subspecies *L. l. lapponica*, *taymyrensis*, and *yamalensis*, the former two of which migrate through previously glaciated western Europe. We also detected unrecognized population structure among bar-tailed godwits wintering in Europe, wherein a significant proportion of purported *lapponica* individuals were in fact *taymyrensis*, necessitating a re-assessment of the migrations, ecology, and population estimates for these subspecies. In the global range of this long-distance migrant, we found evidence of both (1) fidelity to rigid behavioral routines promoting fine-scale geographic population structure (in the east), and (2) flexibility to colonize recently available migratory flyways and non-breeding areas (in the west).

## 1 INTRODUCTION

Animal migration is shaped by, and dependent upon, resources and conditions that vary spatially and seasonally (Alerstam et al., 2003), and therefore migratory populations are considered particularly vulnerable to disruptions caused by climate change and anthropogenic habitat modification (Kubelka et al., 2022; Robinson et al., 2009). In birds, migration involves a highly integrated, multi-trait phenotype (including flight capacity, fueling, navigation, molt, etc.; Piersma et al., 2005) and some degree of endogenous behavioral control (Åkesson et al., 2017; Åkesson & Helm, 2020) that may imply limited flexibility to adapt to conditions outside those in which the migration evolved. Conversely, it is clear that some migratory bird populations have persisted through historical large-scale environmental perturbations, such as repeated Pleistocene glaciations (Batchelor et al., 2019; Hewitt, 2000); apparent responses included population sub-division and size changes, loss and gain of migration, and altered migration routes and distances (e.g., Avise & Walker, 1998; Conklin et al., 2022; Milá et al., 2006; Tan et al., 2023; Zink & Gardner, 2017). A phylogeographic approach (Avise, 2009; Knowles & Maddison, 2002) can help us understand what perturbations migratory populations have survived in the past, and therefore their potential resiliance to future changes.

The bar-tailed godwit, *Limosa lapponica*, is a migratory shorebird (Order Charadriiformes, Family Scolopacidae) that breeds on Arctic and sub-Arctic tundra from Scandinavia east to Alaska, and spends the non-breeding season on intertidal mudflats from West Africa to New Zealand (Figure 1). There are up to six recognized subspecies of bar-tailed godwits, with distinct breeding areas and migratory routes (Figure 1), and further distinguished by variation in plumage and body size (Bom et al., 2022; Engelmoer & Roselaar, 1998; Tomkovich, 2010). Based on non-breeding census data, population sizes for these subspecies vary from c. 100,000 to 500,000 individuals (Wetlands International, 2023), except for the subspecies *L. l. anadyrensis*, which has a poorly described non-breeding range (Chan et al., 2022) but is believed to number <10,000 individuals. The species is considered Near Threatened by the IUCN based on widespread population declines, particularly associated with limitations at migratory stopover sites (Rakhimberdiev et al., 2018; Studds et al., 2017), although the nominate European-wintering population *L. l. lapponica* is thought to be increasing (BirdLife International, 2023).

**Figure 1.**
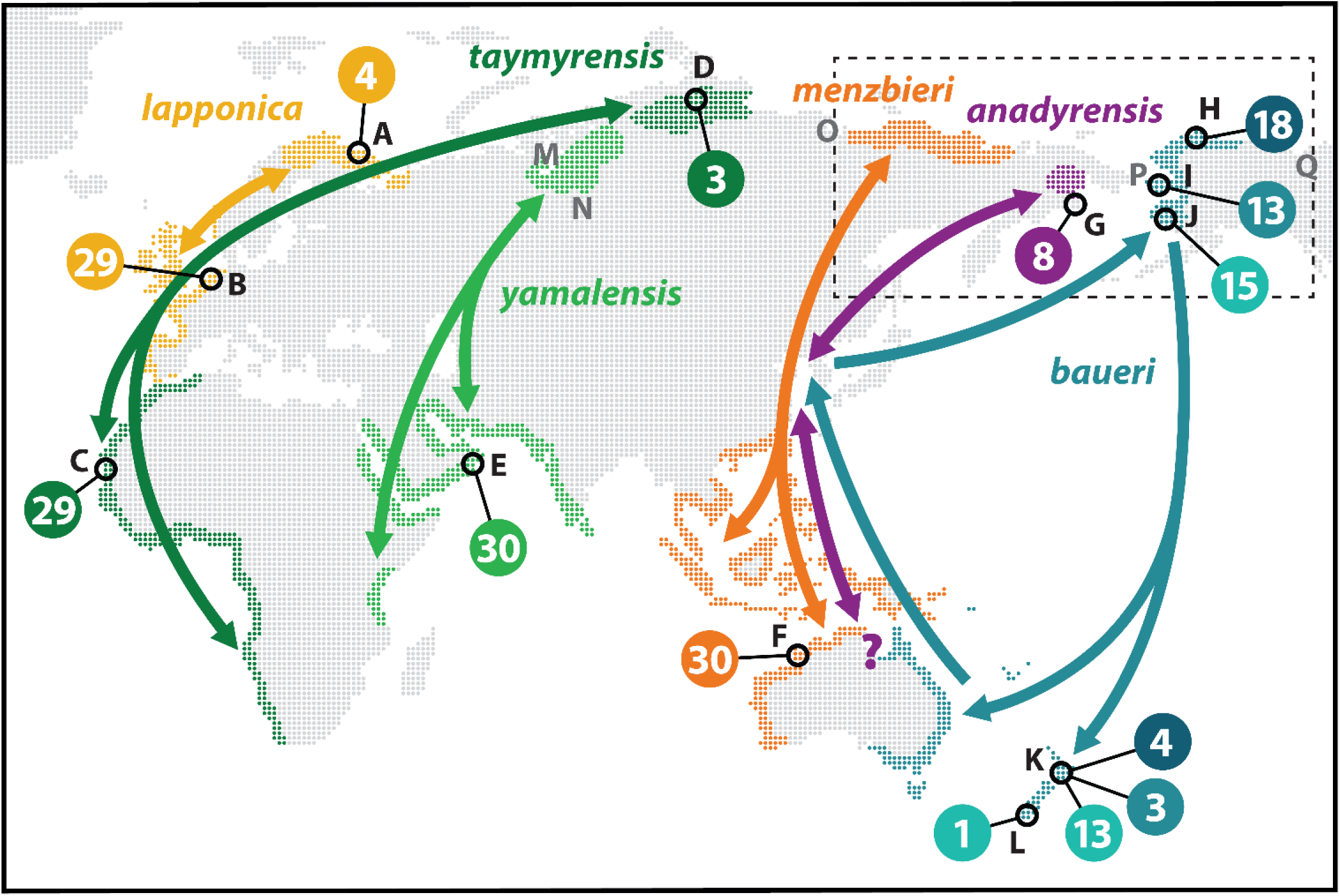
Global distribution and sampling of bar-tailed godwits. For each of six recognized subspecies (indicated by colour), arrows indicate general migration routes between northern breeding areas and southern non-breeding (i.e. boreal winter) areas (coloured areas). Numbers indicate total individuals sampled in each area (black circles); three shades of blue indicate sampling for *baueri* North, Central, and South. Letters refer to sampling sites (in black; see Table S1 for details) and other locations mentioned in the text (in gray): A, Murmansk, Russia; B, Wadden Sea, The Netherlands; C, Banc d’Arguin, Mauritania; D, Taimyr Peninsula, Russia; E, Barr al Hikman, Oman; F, Roebuck Bay, Australia; G, SE Chukotka, Russia; H, Colville River, Alaska; I, Seward Peninsula, Alaska; J, Yukon-Kuskokwim Delta, Alaska; K, North Island, New Zealand; L, South Island, New Zealand; M, West Siberian Lake; N, West Siberian Plain; O, Lena River Delta; P, Bering Land Bridge; Q, Mackenzie River Delta. Dashed box indicates approximate extent of the Beringia region.

In the western Palearctic, the present-day ranges of three bar-tailed godwit subspecies (*lapponica*, *taymyrensis*, and *yamalensis*) were strongly affected by the glaciation of Europe during the last glacial maximum (LGM, c. 20,000–25,000 years before present, ybp; Hewitt, 2000). Accordingly, these three subspecies were indistinguishable at the mtDNA Control Region (Bom et al., 2022), suggesting a recent postglacial radiation into three migration routes (Figure 1). In particular, the entire present-day breeding range of *lapponica*, which makes the shortest migration in the species (one-way distance of c. 2,500–4,500 km), was unavailable at the LGM, making it presumably the youngest subspecies.

By contrast, the eastern part of the bar-tailed godwit breeding range falls within Beringia, an area of the Arctic that stretches from the Lena River in northeast Russia to the Mackenzie River in northwest Canada (Figure 1) and was largely ice-free at the LGM (Ehlers & Gibbard, 2007; Pielou, 1991) and previous glacial cycles (Batchelor et al., 2019). The much deeper biogeographical history of this region, which includes the periodic appearance and disappearance of the Bering Land Bridge, connecting North America with Asia, is reflected in complex histories of diversification in many Beringian taxa (Winker et al., 2023). Thus, two eastern subspecies (*menzbieri* and *baueri*) could have persisted near their present-day ranges for much longer than the western subspecies, and indeed show deeper divergences at the mitochondrial control region (mtDNA) (Bom et al., 2022). A latitudinal cline in both morphology and annual-schedule timing across Alaska (Conklin et al., 2010, 2011) suggests that additional population genetic structure may exist within *baueri*. Conversely, the small breeding range of *anadyrensis* in Chukotka (Figure 1) was at least partly glaciated at the LGM (Batchelor et al., 2019), and therefore could represent a recent expansion into newly available breeding habitat.

Of particular interest is the evolution and persistence of the migration of the *baueri* subspecies, which includes three separate trans-oceanic flights of >5,000 km, including a southward flight of c. 12,000 km across the Pacific Ocean from Alaska to New Zealand (Figure 1), the longest non-stop flight recorded in any landbird (Conklin et al., 2017; Gill et al., 2009). Given this annual routine’s apparent reliance on the specific distribution of habitats and environmental conditions around the Pacific (Conklin et al., 2017; Gill et al., 2014; Piersma et al., 2022), it is unclear how long godwits have been making this migration, and whether it will withstand major changes (e.g. in habitat, prevailing winds etc.) in the future. Unraveling the history of bar-tailed godwits may help reveal whether the trans-Pacific migratory route, and the physiological capabilities required to use it, arose *in situ*, or were products of eastward expansion from the Palearctic (Hedenström, 2010; Piersma et al., 2022).

In this study, we describe the history of divergences and degree of neutral genetic differentiation among global flyway populations of bar-tailed godwits. We used nextera-tagmented, reductively-amplified DNA (nextRAD) sequencing (Russello et al., 2015) for *de novo* discovery of genome-wide single-nucleotide polymorphisms (SNPs) for population genetic analyses, and compared hypothesized evolutionary scenarios in an Approximate Bayesian Computation (ABC) framework using DIYABC-RF (Collin et al., 2021). By reconstructing the recent evolutionary history of bar-tailed godwits, we (1) identify previously unrecognized sub-specific structure, (2) reveal the divergence times and historical connections among populations, and (3) provide insights regarding the origins and maintenance of population structure in long-distance migratory birds.

## 2 MATERIALS AND METHODS

### 2.1 Sampling and DNA extraction

We assembled DNA samples representing all recognized and hypothesized breeding populations within the global range of bar-tailed godwits (Figure 1, Table S1). Where possible, we used samples collected in known breeding areas; however, because bar-tailed godwits breed in low densities in remote regions, for some populations there were few or no breeding samples available. In these cases, we used samples collected from non-breeding sites when sampled individuals could be confidently assigned to breeding areas either because they were remotely tracked to breeding areas using satellite-telemetry or light-level geolocation, or because long-term research programs had established strong links between breeding and non-breeding areas (e.g. through mark-recapture/resight programs). There were no verifiable samples available from the poorly-studied *anadyrensis* population; however, godwits have been collected from coastal southeast Chukotka, within 300 km of the known *anadyrensis* breeding range (Tomkovich, 2010) and far from the expected migration routes of *menzbieri* and *baueri* (Figure 1). Therefore, we included eight such samples, in order to detect genetic variation that might indicate the distinctiveness of *anadyrensis*. All tissue samples were acquired from museum collections or collected by the authors and colleagues in the field under all requisite permits appropriate to their respective countries and institutions.

We extracted genomic DNA from samples using two methods. For blood or organ tissue samples preserved in 95% ethanol, we used the DNeasy Blood and Tissue Kit (Qiagen) following the manufacturer’s instructions for tissue. For blood samples preserved in Queen’s lysis buffer, we used the NucleoSpin Blood QuickPure Kit (Macherey-Nagel). Extract quality was first assessed on a 1.5% agarose gel to exclude extractions with insufficient yield or excessively degraded DNA. We then quantified DNA concentrations using a Qubit 3.0 fluorometer (Life Technologies), diluted extracts to achieve relatively even concentrations, and dried down samples in a SpeedVac concentrator (ThermoFisher Scientific). We delivered 80–162 ng of DNA of 209 individual godwits (Table S1) for SNP discovery and genotyping.

### 2.2 SNP genotyping using nextRAD sequencing

Genomic DNA was converted into nextRAD genomic fragment libraries (SNPsaurus, LLC, USA) following the method described by Russello et al. (2015). Genomic DNA was first fragmented with Nextera reagent (Illumina, Inc.), which also ligates short adapter sequences to the ends of the fragments. The Nextera reaction was scaled for fragmenting 10 ng of genomic DNA, although 20–30 ng of genomic DNA was used for input to compensate for degraded DNA in the samples and to increase fragment sizes. Fragmented DNA was then amplified, with one of the primers matching the adapter and extending 10 nucleotides into the genomic DNA with the selective sequence ‘GTGTAGAGCC’. Thus, only fragments starting with a sequence that can be hybridized by the selective sequence of the primer were efficiently amplified. PCR amplification was done at 74°C for 27 cycles. The nextRAD libraries were sequenced on an Illumina HiSeq-4000 at the Genomics Core Facility, University of Oregon, USA.

For SNP-calling, reads were first trimmed using custom scripts (SNPsaurus, LLC) in bbduk (BBMap tools; Bushnell, 2016). Next, a *de novo* reference was created from abundant reads (after removal of low-quality (phred-scale quality <20) and very high-abundance reads) and reads that aligned to these. All 205,854,244 reads were mapped to the reference with an alignment identity threshold of 95% using bbmap (BBMap tools). Genotype calling was performed using SAMtools and BCFtools (samtools mpileup -gu -Q 10 -t DP, DPR -f ref.fasta -b samples.txt | bcftools call -cv - > genotypes.vcf), applying a minimum read depth filter of 7x. The genotype table was then filtered using VCFtools v.01.14 (Danecek et al., 2011) to remove SNPs called in <80% of samples and putative alleles with a population frequency <3%, to exclude artefactual variants. The resulting VCF file included 6,424 unique loci (150-bp sequences) containing 21,031 SNPs.

Additional cleaning was performed using VCFtools to remove indels (*n* = 478 SNPs), low-quality SNPs (phred-scale quality [QUAL] <999; *n* = 15 SNPs), and samples that failed to sequence (*n* = 1 *menzbieri* individual) or were of unknown breeding population (*n* = 9 individuals from New Zealand that could not be confidently assigned to a breeding region within Alaska based on tracking data). Next, SNPs deviating from Hardy-Weinberg (H-W) proportions in all seven hypothesized populations (excluding *anadyrensis*) were identified and removed using VCFtools (*p* < 0.05; *n* = 48 SNPs); the small sample size in *anadyrensis* made it statistically insensitive to H-W deviations. To ensure independence of loci, we used PLINK v.1.9 (Chang et al., 2015) to identify and remove all SNPs in linkage disequilibrium (LD; *r*^2^ >0.20; *n* = 6,188 SNPs). The LD-pruned data set included 199 individuals and 14,318 unlinked SNPs on 6,140 loci.

We then used VCFtools to calculate proportion of missing SNP calls per individual (range 1– 77%) and removed 15 individuals with >30% missing data. Because inclusion of related individuals can bias population genetic analyses (Anderson & Dunham, 2008; Rodríguez-Ramilo & Wang, 2012), we estimated individual pairwise relatedness in PLINK using the Identity-by-Descent estimator PI HAT; we found no cases of relatedness involving half-siblings or closer (PI HAT >0.20). The final data set included 184 individuals and 14,318 SNPs. We used VCFtools, PLINK, and PGDspider v.2.1.0.3 (Lischer & Excoffier, 2012) to convert data to different formats required for further analysis.

### 2.3 Inference of population structure and diversity

To assess major axes of genetic variation and structure, we performed a principal component analysis (PCA; Patterson et al. 2006) using the R packages *gdsfmt* v.1.14.1 and *SNPRelate* v.1.12.2 (Zheng et al., 2012). We determined the number of ancestral populations or clusters (*K*) and ancestry proportions of these for each individual using ADMIXTURE v.1.3.0 (Alexander et al., 2009). We conducted 20 replicate runs (with random seeds) for each putative number of clusters (*K*) ranging from 1 to 8. We used cross-validation (CV) (Alexander & Lange, 2011) to assess how the CV error rate varied with increasing *K* values and which value provided the lowest CV error. Then, we used the CLUMPAK (Cluster Markov Packager Across K; Kopelman et al., 2015) web server (http://clumpak.tau.ac.il/) with default settings to summarize estimates of individual ancestry proportions to each cluster across replicate runs, inspect potential distinct solutions identified, and visualize the most likely ancestry proportions at each value of *K*.

Globally and for each putative population, we characterized genetic diversity by calculating nucleotide diversity (π), heterozygosity, and inbreeding coefficient (*F*_IS_) using VCFtools. We evaluated evidence for deviations from mutation-drift equilibrium, indicative of departures from a null hypothesis of constant population size, by estimating the per-locus Tajima’s *D* values for each population using VCFtools. We estimated the degree of genetic differentiation among populations by calculating pairwise *F*_ST_ (Weir & Cockerham, 1984) using the *diffCalc* function in the R package *diversity* v.1.9.90 (Keenan et al., 2013), with 95% confidence intervals derived from 500 bootstraps. We calculated *p*-values for *F*_ST_ estimates using the *pairwiseTest* function (1,000 permutations) in the R package *strataG* v.2.4.905 (Archer et al., 2017).

### 2.4 Population evolutionary relationships and demographic history

To visualize evolutionary relationships among populations, we first constructed a midpoint-rooted neighbor-joining tree based on Nei’s genetic distance (Nei, 1972) using the R packages *poppr* v.2.8.5 (Kamvar et al., 2014) and *ape* v.5.3 (Paradis & Schliep, 2019), with missing data replaced by mean allele counts, and node support calculated from 1,000 bootstraps.

To identify likely potential sources of admixture in the tree, we first conducted the three-population test for admixture (Reich et al., 2009) in R using ADMIXTOOLS 2.0.0 (Maier et al., 2023) for all possible population triads. Briefly, the *f*_3_ statistic (Patterson et al., 2012) assesses shared genetic drift to test the null hypothesis that a target population (A) is related to two other populations (B, C) in a simple tree-like manner. We tested significance of the *f*_3_ test using a blocked-jackknife resampling of SNPs, considering blocks of 100 SNPs. A significantly negative value of *f*_3_ (A;(B,C)) indicates that the target population must have arisen from an admixture of the other two populations, or populations closely related to them.

To further investigate the genetic relationships among godwit populations, we estimated population networks using the approach implemented in ADMIXTOOLS 2.0.0 (Maier et al., 2023), which uses the shared and private genetic ancestry components of genome-wide allele frequency data to infer population branching while accounting for potential historical migration and admixture events among populations. To find the best network topology, we used the function *find graphs* for 0 to 5 admixture events (*m*), each with 100 replicates and a maximum of 300 generations. For each replicate, we calculated the out-of-sample score to account for the difference in likelihood score caused by an increasing number of admixture events, which also increase the degrees of freedom. We used the *baueri* South population as a root to fold the networks; even if that population is not an actual outgroup, it allowed visualizing population relationships in a consistent way. The actual population branching order was later assessed statistically using the Approximate Bayesian Computation (ABC) (Beaumont et al., 2002) Random Forest (RF) statistical framework (Pudlo et al., 2016; Raynal et al., 2019). For the best-supported replicates (i.e., those displaying the likelihood score closest to zero), we assessed the goodness-of-fit with the R package *admixture-graph* v.1.0.2 (Leppälä et al., 2017). This approach allows comparing the observed and expected values of *f*_4_ statistics among the different alternatives and identifying the graph that best fits the data (i.e., the one properly predicting the observed *f_4_* values for all possible four-population combinations).

Finally, we evaluated the support for different possible scenarios of population divergence and admixture using the ABC-RF statistical framework (Pudlo et al., 2016; Raynal et al., 2019) in the R package DIYABC-RF v.1.2.1 (Collin et al., 2021). ABC-RF can estimate posterior probabilities of historical scenarios, based on massive coalescent simulations of genetic data. Simulations are compared to observed SNP data using summary statistics to identify the best-fitting model by calculating the number of RF votes and to approximate the posterior probability for the best model (Pudlo et al., 2016). The best-fitting posterior parameter distribution values for the best model can be estimated using a RF procedure applied in a regression setting (Raynal et al., 2019). Estimated parameters include the effective size (*N_e_*) for each population, admixture events and their rates (*r*), and split and admixture times (*t*). We evaluated potential evolutionary scenarios in a stepwise manner, as follows.

As our primary question arising from the above analyses concerned the nature and relative timing of the admixed origin of *menzbieri* (see *Results*), we sought to first reduce the number of potential scenarios by fixing the topology of the relationships among the three *baueri* populations. So, in Step 1, we tested whether the apparently clinal variation among these populations was best represented by one of two possible simple branching topologies or with *baueri* Central as an admixture of *baueri* North and South (Figure S4, Table S2).

In Step 2, we created four possible scenarios (Figure S5c–f) for an admixed origin of *menzbieri* from neighboring populations to the west and east, while allowing populations on the two main branches to diverge at different times relative to each other. Essentially, the admixed origin of *menzbieri* could occur: 1) before either branch began to diverge (Figure S5c); 2) after populations on one branch diverged but before the other branch began to diverge (Figure S5d,e); or 3) after populations on both branches had diverged (Figure S5f). To compete with these scenarios, we created two “null-hypothesis” scenarios in which *menzbieri* was not admixed, but simply diverged from one of the two main branches (Figure S5a,b).

The scenario parameters were considered as random variables drawn from prior distributions (Tables S2 and S3). We used DIYABC-RF to simulate 20,000 genetic data sets per scenario, with the same properties as the observed data set (number of loci and proportion of missing data). Simulated and observed data sets were summarized using the whole set of summary statistics proposed by DIYABC-RF for SNP markers (Table S4), describing population genetic variation (e.g. heterozygosity, proportion of monomorphic loci), differentiation (e.g. *F*_ST_ and Nei’s genetic distances), or admixture (e.g. admixture coefficient, *f_3_* and *f_4_* statistics; Patterson et al., 2012); see the full list and details in Table S4. Linear discriminant analysis (LDA) components were also used as additional summary statistics (Estoup et al., 2012). The total number of summary statistics was 50 and 1,708 for Step 1 and 2, respectively.

We used the random forest (RF) classification procedure (Pudlo et al., 2016) to compare the likelihood of the competing scenarios at each step with DIYABC-RF. RF is a machine-learning algorithm that uses hundreds of bootstrapped decision trees to perform classification, using the summary statistics as a set of predictor variables. Some simulations are not used in decision tree building at each bootstrap (i.e., the out-of-bag simulations), and are used to compute the “prior error rate,” which provides a direct method for estimating the cross-validation error rate. At each step, we built a training set of 20,000 simulated data sets per scenario, with the same number of loci and individuals as the observed data set, and then grew classification forests of increasing size (100, 500, and 2,000 trees) to test convergence. The RF computation provides a classification vote for each scenario (i.e., the number of times a model is selected from the decision trees). We selected the scenario with the highest classification vote as the most likely scenario, and we estimated its posterior probability following the recommendation of Pudlo et al. (2016). We assessed the global performance of our chosen ABC-RF scenario, by calculating the prior error rate based on the available out-of-bag simulations and we repeated the RF analysis (3 times in Step 1; 10 times in Step 2) to ensure that the results converged.

For the best-supported model in Step 2, we estimated posterior distribution values of all parameters using a regression by RF methodology (Raynal et al., 2019) in DIYABC-RF. We used a classification forest of 500 decision trees, based on a training set of 100,000 simulations, to conduct the parameter estimations. We converted divergence time estimates to years assuming a generation time of 8 years (delayed maturity with adult annual survival c. 0.86; Méndez et al., 2018) and the genome-wide mutation rate calculated by Zhang et al. (2014) for Order Charadriiformes: 1.5 x 10^-9^ substitutions per site per year. All of the steps involved in the ABC-RF analysis (simulations, computation of summary statistics, model checking, scenario comparisons, and estimations of parameter posterior distributions) were performed with DIYABC-RF.

## 3 RESULTS

### 3.1 Summary of nextRAD SNP data set

The final data set comprised 184 unrelated bar-tailed godwits genotyped at 14,318 unlinked high-quality SNPs; this included 15–31 individuals in each of seven hypothesized populations, but only three purported *anadyrensis* individuals (Table S1). On average, individuals were genotyped at 93.7% (range: 72–99%) of SNPs and with a mean read depth of 50.4 (range: 15–116). Each SNP was genotyped in an average of 172.5 (range: 149–184, or 81–100%) individuals. Globally, nucleotide diversity (π) was 0.245, and heterozygosity was 20.6%.

### 3.2 Population structure and diversity

The first three axes (PC1–3) of the principal component analysis explained 4.0% of the total genetic variation (Figure 2a). On PC1, individuals were essentially distributed on an axis of longitude in three clusters: a western Palearctic group containing all of *lapponica, taymyrensis* and *yamalensis*; an Alaska group containing all of *baueri*; and an intermediate group of *menzbieri*, suggesting an admixed origin (Figure 2a). PC2 separated 12 *lapponica* individuals from the main western Palearctic cluster. PC3 stretched *baueri* individuals along a latitudinal cline, partly separating the North, South and Central populations. The three purported *anadyrensis* individuals did not form a coherent cluster, one clustering with *menzbieri* and two with *baueri*. Higher-order axes of variation (PC4–6) explained an additional 2.2% of variation but revealed negligible further sub-structure (Figure S1).

**Figure 2.**
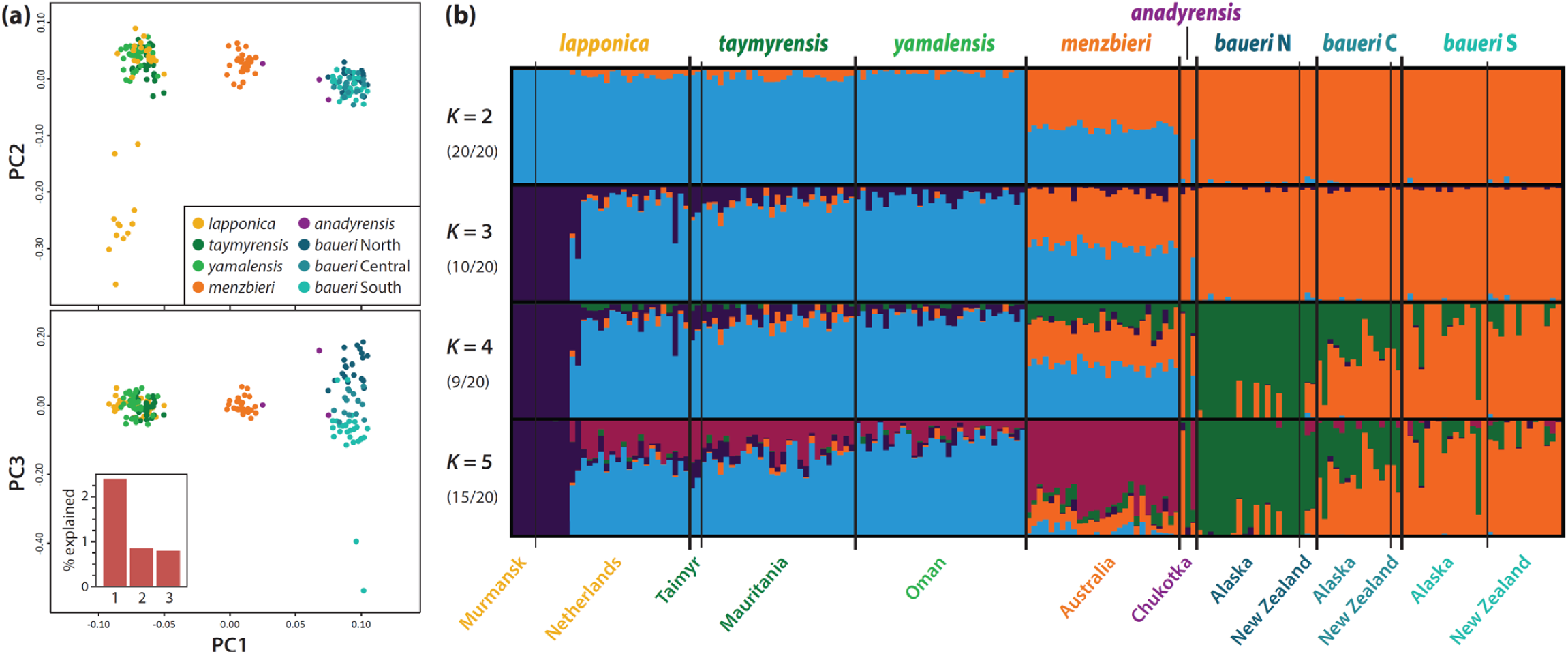
(a) Population structure estimated by principal component analysis. The individual scores (*n* = 184 individuals) for the first three principal components (PC1–3) are shown. The scree plot (inset) indicates the proportion of explained genetic variance by each PC, with PC1–3 explaining 4.0% of total variation. (b) Individual genetic ancestries assigned to major clusters for *K* = 2–5 estimated using ADMIXTURE. At each value of *K*, the ancestry proportions for the 184 individuals for the dominant solution were determined by CLUMPAK summary of 20 replicate runs. Numbers in parenthesis indicate proportion of replicate runs contributing to the dominant solution (see Figure S2 for minor clusters for *K* = 3–5). Names below plot indicate sampling locations (see Figure 1 and Table S1); colors indicate *a priori* purported breeding populations.

Genetic ancestry analyses with ADMIXTURE for *K* = 2 ancestral populations provided the lowest CV error criterion (Figure S2) and 100% support for the major clusters across 20 replicate runs. At *K* = 2, populations were clearly divided between a western Palearctic cluster and an Alaska cluster, with *menzbieri* reflecting admixture between the two (Figure 2b). Additional structure identified in the PCA was apparent at *K* = 3 and 4; i.e. the clear structure within purported *lapponica* and the clinal variation within *baueri*, respectively. At *K* = 5, the identification of a predominantly *menzbieri* cluster may suggest significant genetic drift specific to that population, as opposed to an origin of simple and recent admixture.

Higher values of *K* revealed no further important geographic structure (Figure S2). Again, the purported *anadyrensis* individuals did not form a coherent cluster, but appears to include one *menzbieri* and two *baueri* individuals (Figure 2b).

In both PCA and ADMIXTURE, only 6 of 27 purported *lapponica* individuals sampled on wintering grounds in The Netherlands clearly grouped with the four individuals sampled at known *lapponica* breeding areas in western Russia (see Figure 1, Table S1). Of the remaining 21, 19 clearly grouped with *taymyrensis* (sampled at breeding areas in central Russia and a non-breeding area in Mauritania), and two appear to reflect admixture between *lapponica* and *taymyrensis* (Figure 2a,b). Therefore, for further analysis, we considered a new population of ‘*taymyrensis* Europe’ (*n* = 19), and a reduced *lapponica* population (*n* = 10 ‘true’ *lapponica* plus the 2 admixed individuals). For clarity, we hereafter refer to the original *taymyrensis* samples as ‘*taymyrensis* Africa’ (*n* = 29). As PCA and ADMIXTURE results do not support an identifiable *anadyrensis* population, we excluded those samples (*n* = 3) from further analysis. Therefore, we proceeded with 181 individuals in eight populations.

Globally, nucleotide diversity was 0.245, and this was similar for all populations (Table S5). Mean heterozygosity was also generally uniform (range of means: 0.198–0.215), with no evidence of inbreeding in any population (all mean *F*_IS_ < 0.20; Table S5). Values of Tajima’s *D* were low and similar among populations (range of means: 0.214–0.339; Table S5), suggesting no evidence of dramatic population-size changes or additional hidden structure.

Pairwise *F*_ST_ among the eight hypothesized populations ranged 0.001–0.048 (Table 1), and were greatest between the westernmost and easternmost breeding populations (*lapponica* vs. *baueri; F*_ST_ values ranging from 0.046 to 0.048 for the three *baueri* sub-populations). The two groups that winter in Europe, all previously considered *lapponica* (*lapponica* and *taymyrensis* Europe), were moderately differentiated (*F*_ST_ = 0.016). Differentiation among the three sub-populations of *baueri* (*F*_ST_ = 0.006–0.009) was lower than for most recognized subspecies pairs, with 95% CIs that included zero. The lowest values were among the three groups breeding in central Russia (*taymyrensis* Europe, *taymyrensis* Africa, and *yamalensis*; *F*_ST_ = 0.001–0.003).

**Table 1.**
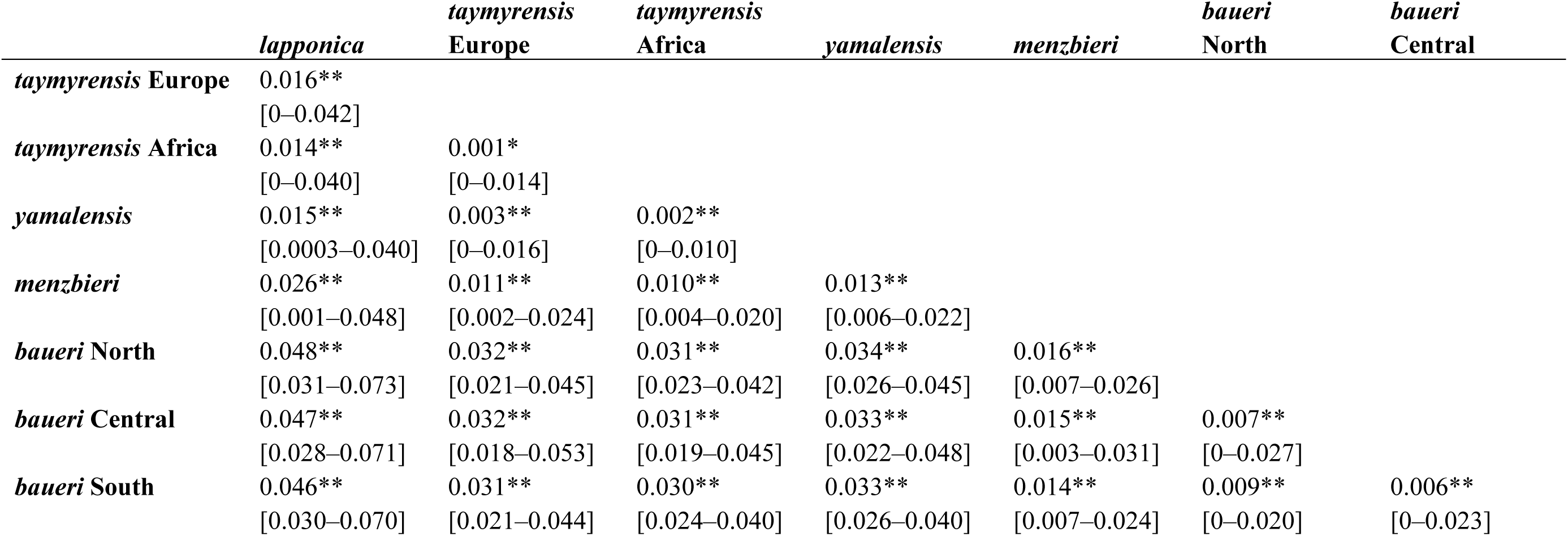
Mean population pairwise *F*_ST_ and 95% confidence intervals of estimates. All comparisons are significantly different from zero (1,000 permutations; *\*p* = 0.038, *\*\*p* < 0.001).

|  | *lapponica* | *taymyrensis*<br>Europe | *taymyrensis*<br>Africa | *yamalensis* | *menzbieri* | *baueri*<br>North | *baueri*<br>Central |
| --- | --- | --- | --- | --- | --- | --- | --- |
| *taymyrensis* Europe | 0.016**<br>[0–0.042] |  |  |  |  |  |  |
| *taymyrensis* Africa | 0.014**<br>[0–0.040] | 0.001*<br>[0–0.014] |  |  |  |  |  |
| *yamalensis* | 0.015**<br>[0.0003–0.040] | 0.003**<br>[0–0.016] | 0.002**<br>[0–0.010] |  |  |  |  |
| *menzbieri* | 0.026**<br>[0.001–0.048] | 0.011**<br>[0.002–0.024] | 0.010**<br>[0.004–0.020] | 0.013**<br>[0.006–0.022] |  |  |  |
| *baueri* North | 0.048**<br>[0.031–0.073] | 0.032**<br>[0.021–0.045] | 0.031**<br>[0.023–0.042] | 0.034**<br>[0.026–0.045] | 0.016**<br>[0.007–0.026] |  |  |
| *baueri* Central | 0.047**<br>[0.028–0.071] | 0.032**<br>[0.018–0.053] | 0.031**<br>[0.019–0.045] | 0.033**<br>[0.022–0.048] | 0.015**<br>[0.003–0.031] | 0.007**<br>[0–0.027] |  |
| *baueri* South | 0.046**<br>[0.030–0.070] | 0.031**<br>[0.021–0.044] | 0.030**<br>[0.024–0.040] | 0.033**<br>[0.026–0.040] | 0.014**<br>[0.007–0.024] | 0.009**<br>[0–0.020] | 0.006**<br>[0–0.023] |

### 3.3 Population evolutionary relationships and demographic history

The neighbor-joining (NJ) tree based on Nei’s distance (Figure 3a) again identified two main branches (western Palearctic and Alaska groups), with low node support within each group, and *menzbieri* occupying an intermediate position between them.

**Figure 3.**
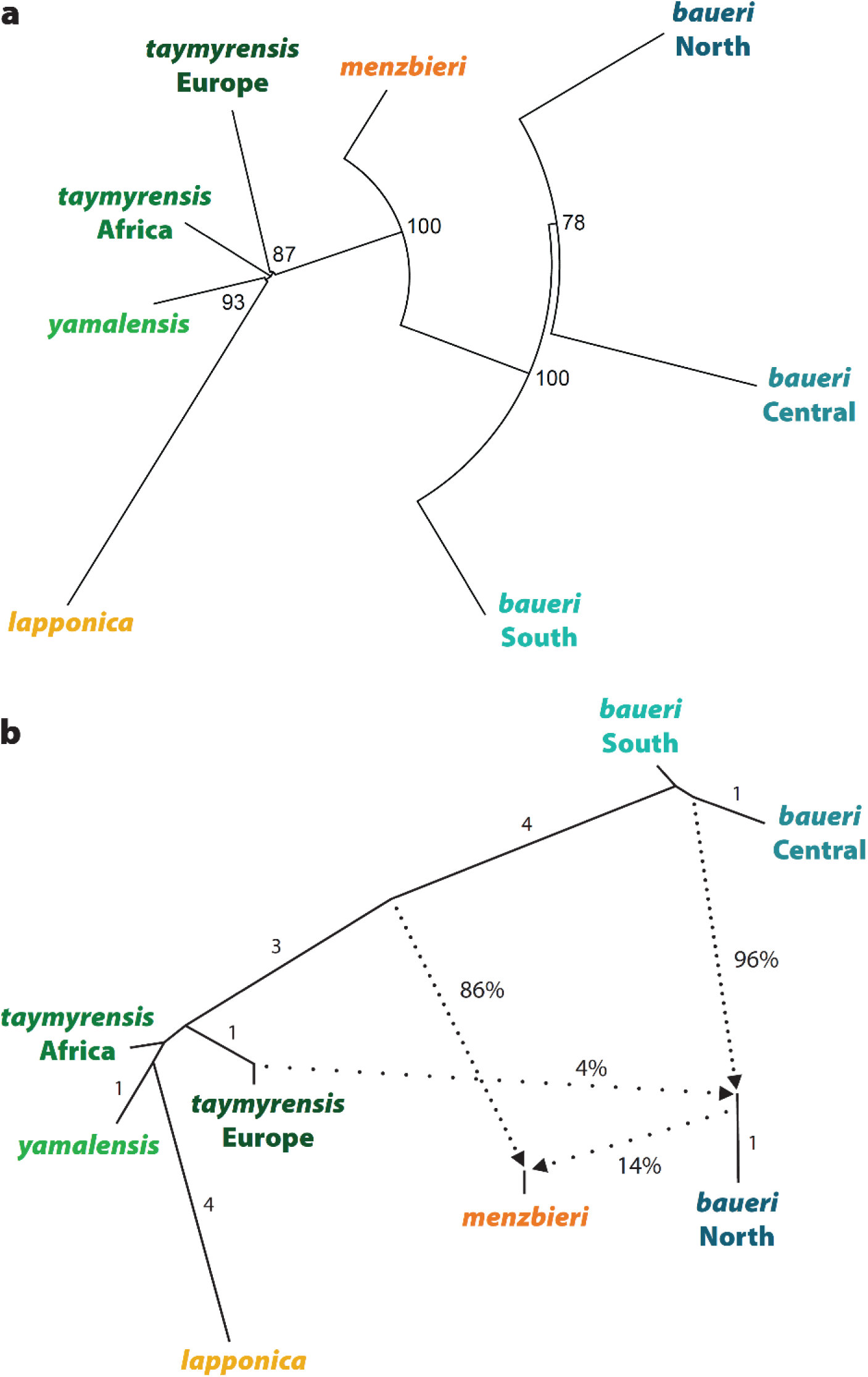
(a) Unrooted neighbour-joining tree, based on Nei’s minimum distance. Numbers indicate bootstrapped node confidence (%). (b) AdmixtureGraph showing best graph with two migration reticulations (*m* = 2), arbitrarily rooted with *baueri* South population (see Methods). Numbers indicate relative genetic distances (unlabeled branches = 0 distance). Dotted lines represent admixtures, with percentages indicating admixture proportions, and arrows indicating direction of admixture.

The three-population test (based on the Patterson’s *f*_3_ statistic of the form (A;(B,C)); Table S6) identified *menzbieri* as a likely product of admixture between the two main branches identified by the NJ tree (see Figure 3a). Of 168 possible population triads, 12 produced a significantly negative *f*_3_ statistic; all of these included *menzbieri* as the target population (A) and one member of each of the two main branches (*lapponica/taymyrensis/yamalensis* or the *baueri* group, respectively) as the source populations (B, C) (Table S6).

For admixture graphs estimated by ADMIXTOOLS 2, the greatest increase in likelihood occurred at *m* = 2 admixture edges in the population graph (Figure S3a), but out-of-sample scores indicated the highest likelihood at *m* = 3 (Figure S3b). Overall, the five best-supported replicates (i.e., those with likelihood scores closest to zero) included two distinct network topologies for *m* = 3 and one for *m* = 2. The best-supported topologies at *m* = 2 and 3 (Figure S3c,d) each provided a perfect fit with the data as estimated by *Admixture-graph* (i.e., no outlying observed *f*_4_ statistic values compared to those theoretically expected from these graphs; Figure S3e,f). Therefore, according to parsimony we present the model at *m* = 2 (Figure 3b). In this graph, *menzbieri* arose from the admixture of a population ancestral to present-day western Palearctic populations (86% contribution) and a population closely related to *baueri* North (14%). Additionally, *baueri* North has received 4% ancestry from a population closely related to *taymyrensis* Europe.

To resolve the nature and relative timing of the admixed origin of *menzbieri* suggested by all previous analyses, we compared the likelihood of alternative scenarios using the approximate Bayesian computation – random forest approach (ABC-RF). In Step 1, we tested whether the apparently clinal variation among the three *baueri* populations was best represented by a simple branching topology or by historical admixture (Figure S4). The best-fitting scenario, representing *baueri* Central as an admixture of *baueri* North and South, received the greatest proportion of RF votes (44.5%), with a posterior probability of 51.0% (Figure S4 and Table S7). This ABC-RF analysis had good power to discriminate among the three scenarios, as suggested by the low prior error rate of 11.4% and the confusion matrix (Table S8).

Therefore, we fixed the topology of the main Alaskan branch accordingly in the global scenarios evaluated in Step 2.

In Step 2 (Figure S5), we compared six possible scenarios for the origin of *menzbieri*: two “null-hypothesis” scenarios in which *menzbieri* simply diverged from one of the two main branches (*a*,*b*), and four in which an admixed *menzbieri* arose at different time points relative to the diversification within each branch (*c*–*f*). The best-supported scenario was *c*, in which the historical admixture occurred prior to the main diversifications within both the western Palearctic and Alaska groups (Figure 4c); this scenario received (mean ± SD) 26.4 ± 0.9% of votes, with a posterior probability of 46.1 ± 2.6% (Figure S5, Table S9). The global prior error rate of 22.0 ± 0.0% indicated that discriminating power among the six scenarios was good, but with some difficulties in discriminating between scenarios *c* and *e* (Table S10). Scenario *e* was the second best-supported scenario, receiving 25.3 ± 1.0% of votes. This scenario was most similar to *c*, but with the admixture occurring before diversification among *baueri* populations and after *lapponica* diverged from the other western Palearctic populations (Figure 4e).

**Figure 4.**
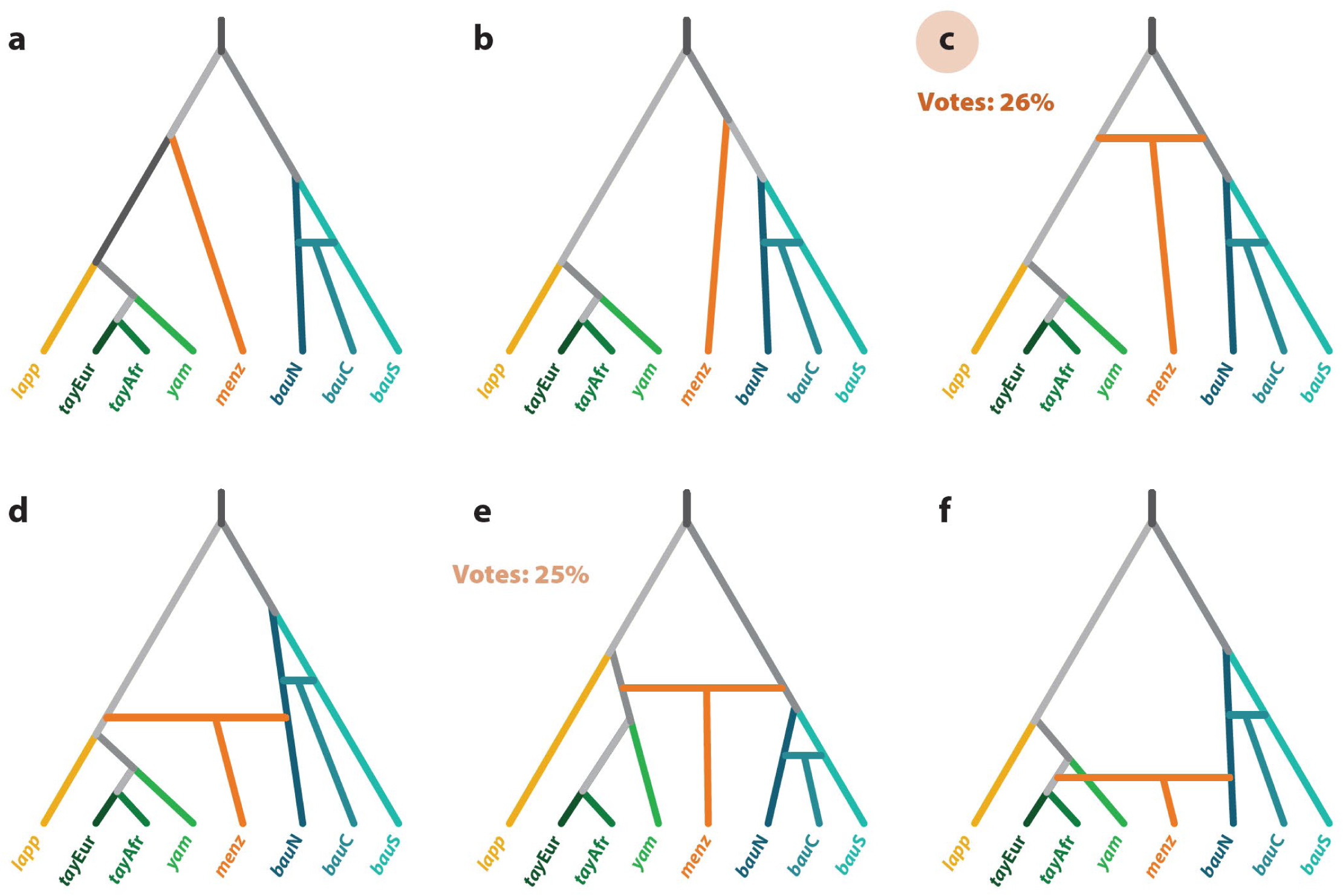
Scenarios tested in Step 2 of DIYABC-RF analysis: six possible scenarios for the origin of *menzbieri*, by either simple branching (a,b) or by admixture of the two main branches at relatively ancient (c), intermediate (d,e) or recent (f) time periods. Extant (sampled) populations are indicated by colours; inferred historical populations are shown in shades of grey. The best-supported scenario was (c), followed by (e), as inferred by the proportion of Random Forest classification votes; see Figure S5 and Tables S9 and S10 for details.

Each of the 20 demographic parameters of the best-supported scenario was estimated within the ABC-RF framework (Table S11). Timing parameter estimates support the existence of three main lineages at the LGM, with the divergence of the western Palearctic and Alaskan branches occurring c. 38.2K ybp (modal estimate; 5^th^–95^th^ quantiles: 22.0–93.1K), followed by an admixture giving rise to *menzbieri* c. 22.0K ybp (13.4–52.5K; Figure 5, Table S11).

**Figure 5.**
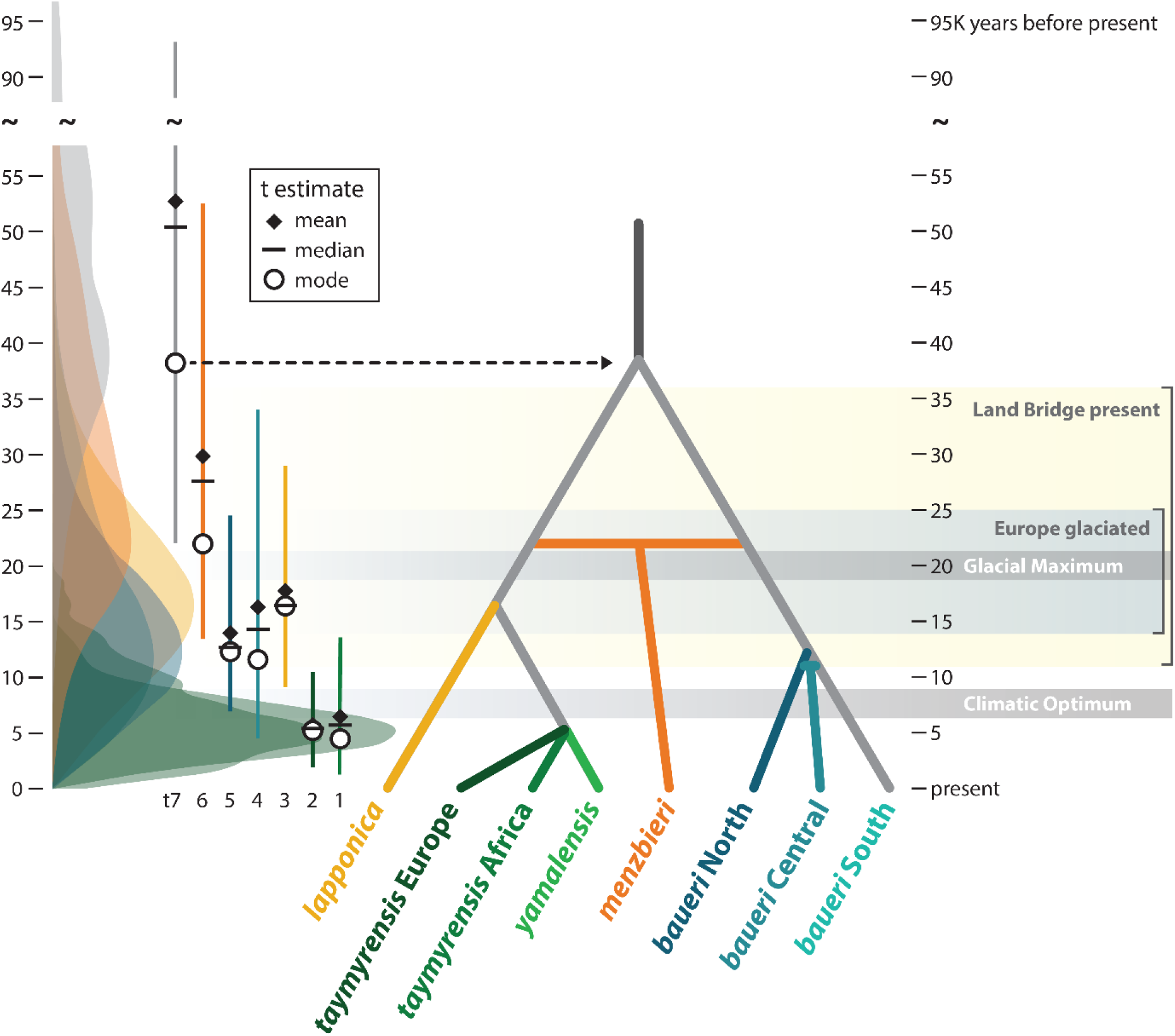
Best-supported Scenario in Step 2 of DIYABC-RF analysis (see Figure 4), scaled to relative time-parameter estimates (converted to years assuming a generation time of 8 years) for five divergence events (branches) and two admixture events (horizontal bars). On left, posterior distributions for each time parameter estimate, indicating mode, mean, median, and interval of 5^th^–95^th^ quantiles (vertical line). For reference, estimated timing of Last Glacial Maximum, Holocene Climatic Optimum, and most recent presence of Bering Land Bridge and glaciation of Europe are shown.

The post-LGM warming period featured *lapponica* diverging from the western Palearctic lineage (c. 16.4K ybp; 9.1–28.9K) and the diversification of *baueri* populations (c. 11.6– 12.3K ybp); although we modeled the *baueri* diversification as a divergence followed by an admixture (see Step 1), the two time estimates were essentially indistinguishable, indicating a relatively contemporaneous radiation of three populations, rather than contact after a prolonged isolation (Figure 5). Most recently, the remaining western Palearctic lineage diversified into *taymyrensis* (Europe and Africa) and *yamalensis* populations (c. 4.5–5.2K ybp); these two divergence times were also indistinguishable, rather than clearly sequential (Figure 5).

## 4 DISCUSSION

### 4.1 Population structure

Previous research showed that the three western Palearctic bar-tailed godwit populations (*lapponica*, *taymyrensis*, and *yamalensis*) were indistinguishable at the mtDNA control region (Bom et al., 2022), consistent with a very recent post-glacial expansion into three breeding areas and migration routes. However, in genome-wide SNPs, we found that *lapponica* was strongly differentiated from *taymyrensis* and *yamalensis*, which were weakly but significantly differentiated from each other (Table 1). This suggests a more complicated phylogeographic history, including a post-LGM colonization by *lapponica* of recently-deglaciated Europe, followed later by emerging structure along two flyways in the source population breeding in the central Russian Arctic. These differences from patterns found in mtDNA illustrate the greater resolution of genome-wide nuclear markers to detect subtle and recent population structure (Edwards & Bensch, 2009).

We also found unexpected structure among samples collected in the Dutch Wadden Sea during winter, purported to belong to *lapponica*: of these 27 samples, only 6 grouped with the 4 individuals sampled in the breeding area of *lapponica*, whereas 19 clearly clustered with *taymyrensis*, and another 2 appeared to represent admixture between the two subspecies (Figure 2). Due to long-term mark-resighting programs on the flyway, we were able to restrict our purported *lapponica* samples to marked individuals that had been observed in multiple years in mid-winter (December–January) in the Wadden Sea and had never been reported in west Africa; this should have provided the best chance to exclude *taymyrensis* individuals from that sample, as *taymyrensis* is expected to pass through western Europe on migration between Russian breeding grounds and non-breeding grounds in west Africa but to be absent in the winter (Figure 1; Drent & Piersma, 1990; Duijns et al., 2012). Our results strongly indicate that both *taymyrensis* and *lapponica* are present in Europe in the winter.

The currently recognized *lapponica* and *taymyrensis* subspecies were once considered differentially migrating populations of the nominate subspecies *L. l. lapponica*, until the recognition of plumage and morphometric differences between the Fennoscandian and central Russian breeding areas led to the full subspecies status of *taymyrensis* (Engelmoer & Roselaar, 1998); this taxon was itself later split with the recognition of *yamalensis* (Bom et al., 2022). *Lapponica* and *taymyrensis* are generally considered ‘leap-frog’ migrants, with the entirely of *taymyrensis* wintering in west Africa, south of European-wintering *lapponica* (Figure 1), consistent with historical mark-resight and tracking evidence (Duijns et al., 2012). By contrast, based on morphometric data, Engelmoer (2008) predicted that approximately 20% of bar-tailed godwits wintering in the Dutch Wadden Sea were *taymyrensis*, and thus the migration along the flyway was in fact a ‘partial leap-frog system’. Despite this uncertainty, the current population estimate for *lapponica* (150,000–180,000 and increasing; Wetlands International, 2023) is based entirely on winter census data from Europe, assuming no mixing of subspecies. If our results from the Wadden Sea (i.e. c. 70% *taymyrensis*) reflect the subspecies proportions of the entire European wintering population, then the currently accepted population estimate for *lapponica* might overstate the actual population size by a factor of three, and any perceived increase could reflect unknown dynamics in two conflated populations. This should be tested through genetic assignment of a larger and geographically broader sample of the European wintering population, followed by a re-evaluation of the population sizes and conservation status of both *lapponica* and *taymyrensis*.

We were unable to completely resolve structure among the three Beringian subspecies due to the small number and high sequencing failure rate of purported *anadyrensis* samples; we thus can conclude little regarding the status or history of that subspecies, which has a distinct migration route through eastern Russia but an incompletely described non-breeding range (Chan et al., 2022; Tomkovich, 2010). However, the three samples in our final data set did not cluster together, grouping with either *menzbieri* (*n* = 1) or *baueri* (*n* = 2; Figure 2). This suggests that these individuals, sampled in southeast Chukotka, were most likely passage migrants from the neighboring subspecies to the west and east, rather than from the closest breeding population, but leaves unanswered the question of the potential genetic distinctiveness of *anadyrensis*. Therefore, genetic evaluation of the status of *anadyrensis* awaits analysis of confirmed breeding members of the subspecies.

Within *baueri*, we confirm that behavioral and morphological differences observed along a latitudinal cline in the Alaska breeding range (Conklin et al., 2010, 2011; Conklin & Battley, 2011) are mirrored by a similar genetic cline (Figure 2). Bar-tailed godwits from the entire Alaska breeding range regularly mix at non-breeding sites in New Zealand (Battley et al., 2020; Conklin et al., 2011) and Australia (Wilson et al., 2007) and at migratory staging sites in the Yellow Sea and southwest Alaska (Battley et al., 2012). In the Yellow Sea, many *baueri* individuals also mix freely in staging flocks of *menzbieri* on northward migration (see Figure 1; Battley et al., 2012; Choi et al., 2015). The clear genetic structure maintained among (sub-) populations that regularly meet in the non-breeding season implies that behavioral rigidity, in terms of strict migration timing and fidelity to a breeding region, is sufficient to preclude panmixia among and within subspecies. Within *baueri*, previous research found no clear link between migration timing, which generally follows the latitudinal genetic cline we detected, and variation at the gene Clock, thought to be associated annual-cycle scheduling in birds (Parody-Merino et al., 2019). Therefore, it remains unclear whether the phenotypic and genetic cline within *baueri* is primarily driven by neutral genetic drift or involves selection, and whether it is emerging or stably maintained over a long time period.

### 4.2 Phylogeographic history

Our demographic reconstruction in DIYABC-RF provides the first estimates for divergence times in bar-tailed godwits, and new insights regarding the historical origins of present-day flyway populations. Our divergence time estimate of c. 38K ybp for the two main lineages suggests the ancestors of the western Palearctic and Alaskan populations existed in Siberia and Beringia well before the LGM (Figure 5). This corresponds to a period (c. 40K ybp) when the Bering Sea separated Asia and North America (Farmer et al., 2023) and much of the Chukotka Peninsula was glaciated (Batchelor et al., 2019); either or both of these factors could have caused or perpetuated isolation of the two lineages. We inferred a subsequent admixture of the two lineages c. 22K ybp, giving rise to present-day *menzbieri* (Figure 5).

During this period, just prior to and including the LGM, Beringia was largely unglaciated (Batchelor et al., 2019) and lower sea levels created a connection between the continents via the Bering Land Bridge (c. 36–11K ybp; Farmer et al., 2023), a situation that could have created new breeding habitat available for colonization from both the west and the east, or conceivably re-connected breeding populations in a continuous swath of tundra habitat stretching from Chukotka to Alaska. Indeed, reconstructed vegetation histories indicate that at least part of the Bering Land bridge hosted graminoid and herbaceous tundra, the preferred breeding habitat of bar-tailed godwits, during the LGM (Wang et al., 2017). Presumably, subsequent re-flooding of the Bering Strait (by c. 11K ybp) and incremental reduction and isolation of tundra habitat associated with the warming period leading to the mid-Holocene Climatic Optimum (c. 8K ybp; Kraaijeveld & Nieboer, 2000; Stewart & Dalén, 2008; Wauchope et al., 2017) served to isolate these three extant lineages into their present-day breeding ranges (Figure 1); similar structure across Beringia is found in numerous avian taxa (Winker et al., 2023)

Because significant portions of Alaska have remained unglaciated for hundreds of thousands of years (Batchelor et al., 2019; Winker et al., 2023) and our results suggest no temporal limit to *baueri*’s occupation of its present-day breeding range, we can infer no theoretical maximum age of that population’s trans-Pacific migration (Gill et al., 2009). Therefore, we cannot help discern between scenarios of *in situ* evolution of these extremely long non-stop flights in the Pacific versus incremental up-scaling of flight distances through eastward expansion of godwits from Asia (Hedenström, 2010; Piersma et al., 2022). We can only infer that any such eastward expansion did not appear to occur recently (e.g. post-LGM), as *baueri* has apparently existed in Alaska for sufficient time for genetic and morphological structure to develop along a latitudinal cline (Conklin et al., 2011). However, our inference of a radiation into this cline c. 12K ybp is potentially consistent with expansion northward or southward within Alaska, or with emerging isolation-by-distance across a stably occupied area.

Paleoclimatic modeling, to assess change in both atmospheric circulation and habitats in the Pacific Basin across time, may reveal more about how long *baueri* could theoretically have been migrating across the Pacific in the current manner (Piersma et al., 2022).

If bar-tailed godwits existed in a Siberian-Beringian refugium (stretching from the West Siberian Plain to Alaska) at the LGM and then expanded westward with the de-glaciation of Europe (c. 14K ybp), then we would expect *lapponica* to have diverged very recently from other western Palearctic lineages, particularly given the expected delay between geographic separation and genetic divergence. In this light, our inferred divergence of *lapponica* c. 16K ybp is unexpected, as their present-day breeding range in Fennoscandia was not available until at least 3,000 years after that (Stroeven et al., 2016). This divergence-time estimate is not easily explained by a simple westward expansion, and favors a scenario of isolation followed by expansion from tundra breeding areas that were available south or southeast of glaciated Europe during or soon after the LGM. In fact, such refugia, including in central/southern Europe, have been inferred for tundra-breeding shorebirds (Arcones et al., 2021) and other taxa (Hewitt, 2004; Sommer & Nadachowski, 2006; Taberlet et al., 1998). If the breeding grounds of *lapponica* gradually moved toward Fennoscandia from central Europe as glaciers retreated north, then its present-day migration route would re-trace that northward expansion, as has been hypothesized for numerous migratory birds that either originally evolved from sedentary populations (Rappole & Jones, 2002; Salewski & Bruderer, 2007) or oscillated between short- and long-distance migration as northern areas were impacted by cyclic glaciations (Bruderer & Salewski, 2008; Zink & Gardner, 2017).

At the LGM, the lineage that recently gave rise to *taymyrensis* and *yamalensis* could have existed east or south of the glaciers that extended from Europe to the Taimyr Peninsula of central Russia (Arcones et al., 2021; Batchelor et al., 2019). We may then expect that the present-day breeding range and migration of *yamalensis* (Figure 1) would most closely resemble the state of the western Palearctic lineage at the LGM. Consistent with this, there is evidence of tundra habitats existing south of the ice-dammed West Siberian Lake at the LGM (Binney et al., 2017; Mangerud et al., 2004), close to the present-day breeding area of *yamalensis* in the West Siberian Plain. Currently, *taymyrensis* and *yamalensis* have some degree of isolation in breeding areas along a latitudinal and longitudinal gradient (Figure 1; Bom et al., 2022). This may suggest that a northeastward expansion of breeding range toward the Taimyr Peninsula was associated with discovery of the migration route of *taymyrensis* through western Europe (Figure 1) after glaciers retreated.

We have presented the first unequivocal evidence that *taymyrensis* can be found in Europe in winter, and it is therefore unclear whether this is a new, perhaps emerging, development in modern times (i.e. tens or hundreds of years) or a stable situation that went previously undetected. However, the detection of two apparent F1 *lapponica*/*taymyrensis* hybrids (see Figure 2) in our relatively small sample from the Wadden Sea appears more indicative of a brief period of significant contact between the subspecies. Also, the negligible differentiation between European- and Africa-wintering *taymyrensis* (*F*_ST_ = 0.001) provides no evidence of further structure within that subspecies. Taken together, our results suggest that the existence of *taymyrensis* in Europe, or at least conditions promoting successful cross-population recruitment, is a recent or increasing phenomenon.

### 4.3 Population structure in long-distance migrants

Highly mobile organisms, such as migratory birds, are interesting cases for studying the effects of spatial isolation on population structure. On one hand, highly mobile species are expected to have high dispersal ability, which should decrease the strength of physical or ecological barriers to gene flow, resulting in shallow structure, or even panmixia, across large geographic areas. Accordingly, a negative relationship between dispersal ability and population structure has been demonstrated in various taxa (Bohonak, 1999; Medina et al., 2018). In birds, both migratory tendency and flight capacity *per se* have been linked with increased dispersal and decreased population structure and speciation (Arguedas & Parker, 2000; Claramunt et al., 2012). By this reasoning, we should expect unimpeded gene flow in an extreme long-distance migrant like the bar-tailed godwit, which can fly distances >11,000 km across formidable ecological barriers without resting, eating, or drinking (Gill et al., 2009; Piersma et al., 2022), predicting negligible sub-specific population genetic structure.

Paradoxically, migration has also been linked with increased rates of speciation and diversification in birds (Rolland et al., 2014; Winger et al., 2014). Long-distance migration is associated with high site-fidelity and rigid behavioral and physiological traits that enable timely exploitation of predictable seasonal resources across hemispheres (Åkesson et al., 2017; Conklin et al., 2013; Winger et al., 2019). These traits predict low effective dispersal among neighboring populations, either through strict spatial or temporal isolation, or by preventing successful recruitment by immigrants (Nosil et al., 2005). In this light, we might expect bar-tailed godwits to exhibit strong structure among flyway populations.

Within this species, we see evidence for both ends of the continuum: in the west, negligible structure between two populations with adjacent breeding areas but separate ranges in the non-breeding season (*taymyrensis* and *yamalensis*), and in the east, clinal structure within a single subspecies, where individuals freely mix throughout the non-breeding season (*baueri*). The latter situation, involving birds making the longest non-stop migratory flights on earth, demonstrates the weak link between dispersal *ability* (in terms of flight capacity) and actual dispersal. In fact, bar-tailed godwits are known for extremely rigid site-fidelity at non-breeding and migratory staging sites (Conklin et al., 2013; Rakhimberdiev et al., 2018), and are presumed to show similar fidelity to breeding sites as adults. The very fine-scale structure we demonstrate within the Alaska breeding range suggests that natal dispersal is also quite restricted in *baueri* (i.e. relative to distances of c. 400–800 km between our sampling sites; Figure 1). As we have no reason to expect weaker site-fidelity in other bar-tailed godwit populations, we presume the weak differentiation between *taymyrensis* and *yamalensis*, which may meet at pre- and post-breeding areas and breed <500 km from each other (Bom et al., 2022), is a result of their recent divergence (c. 6,000 ybp) rather than high levels of gene flow.

While not clearly related to dispersal and gene flow, flight capacity may be more indicative of a species’ propensity for exploration and colonization of new migratory flyways. As illustrated in other long-distance migratory shorebirds, such as dunlin, *Calidris alpina* (Wenink et al., 1996), and red knot, *C. canutus* (Conklin et al., 2022), such species appear to respond relatively quickly to major climatic changes, such as retreat of glacial ice sheets, with establishment of new flyway populations (as we have demonstrated with the *lapponica* subspecies). As migration of adult birds tends to be more conservative, such ‘innovations’ in a population may result largely from inexperienced juveniles conducting their first migration (Cresswell, 2014; Verhoeven et al., 2022). Intriguingly, bar-tailed godwits and other long-distance migrants also seem flexible (at least in evolutionary timescales) with regard to migration distance. For example, *lapponica* and *yamalensis*, among the youngest of bar-tailed godwit subspecies, migrate as little as 2,500–5,000 km between breeding and non-breeding grounds, whereas the other four subspecies travel one-way distances of 10,000–18,000 km (Figure 1). This shows that behavioral and physiological ‘upscaling’ to accommodate the world’s longest flights (Hedenström, 2010) is not a one-way journey; it may be reversed, with regression toward short-distance migration, if the ecological opportunity arises.

## ACKNOWLEDGEMENTS

We thank Per Palsbøll and Ritsert Jansen for guidance and help to develop the funding proposal, and to members of the Palsbøll and Fontaine laboratories at the University of Groningen and Montpellier for helpful discussions during project conception and analysis, respectively. We thank Anneke Bol at NIOZ Royal Netherlands Institute for Sea Research for DNA extractions of Mauritania samples; Marco van der Velde for laboratory assistance in Groningen; Eric Johnson and Paul Etter (SNPsaurus) for sequencing, bioinformatics, and technical expertise; and the Centre for Information Technology of the University of Groningen (particularly Bob Dröge and Cristian Marocico) for technical support and access to the Peregrine high-performance computing cluster. For providing samples, we thank Massey University, Moscow Lomonosov State University Zoological Museum, NIOZ, Royal Ontario Museum (Oliver Haddrath and Mark Peck), U.S. Geological Survey Alaska Science Center, University of Groningen, and University of Washington Burke Museum. For godwit capture and field sampling, we thank Luke DeCicco, Jimmy de Fouw, Maks Dementyev, Nick Hajdukovich, Jim Johnson, Murray Potter, Adrian Riegen, David Melville, Dan Mulcahy, Adrian Riegen, Rob Schuckard, Craig Steed, and Cleland Wallace, plus numerous local volunteers. In Australia, we thank Broome Bird Observatory and the Australasian Wader Studies Group for logistical and field work support, and we acknowledge the Yawuru People via the offices of Nyamba Buru Yawuru Limited for permission to capture godwits on the shores of Roebuck Bay, traditional lands of the Yawuru people. Sampling was supported by the David and Lucile Packard Foundation (Alaska, Australia, and New Zealand), BirdLife Netherlands and WWF-Netherlands (Australia), and a Royal Society of New Zealand Marsden Fund grant to P.F.B. and Andrew Fidler (New Zealand). The Prins Bernhard Cultuurfonds grant to T.P. started up the long-term efforts in Mauritania, supported now by operational grants from NIOZ; we thank the authorities of the Parc National du Banc d’Arguin for access and support. Work in the Wadden Sea was supported by Waddenfonds (project Metawad, WF209925) and operational funding from NIOZ. Work in Oman was financially supported by the Research Council (TRC) of the Sultanate of Oman (ORG/EBR/12/002). The core of this project was supported by a Dutch Research Council (NWO) grant to T.P. (ALW-Open Programme grant, ‘Ecological drivers of global flyway evolution’ 824.01.001). Any use of trade, product or firm names is for descriptive purposes only and does not imply endorsement by the U.S. Government.

## AUTHOR CONTRIBUTIONS

J.R.C., Y.I.V., M.C.F. and T.P. conceived and designed the study. P.F.B., R.A.B., R.E.G., C.J.H., J.t.H., D.R.R., T.L.T., N.W., and P.S.T. organized and performed field sampling and/or maintained individual resight-history databases. J.R.C. and Y.I.V. curated samples and conducted the labwork. J.R.C., M.J.M.L. and M.C.F. analysed and interpreted the data, with assistance from Y.I.V. J.R.C. and M.C.F. wrote the manuscript, with major contributions from Y.I.V. and T.P. All authors provided feedback and helped edit the manuscript.

## DATA AVAILABILITY STATEMENT

Demultiplexed nextRAD short-read data were deposited in NCBI’s SRA archives under BioProject ID xxx (TBA). Associated files (VCF and metadata) were deposited in DRYAD (doi: TBA).

## BENEFIT-SHARING STATEMENT

All genetic samples were collected with collaboration and/or consultation with local communities, and all principal collaborators who provided samples are included as co-authors. The research addresses an international priority concern, which is the conservation of the study organism. Benefits from this research accrue from the sharing of our data and results with the scientific and stakeholder communities, both directly and on public databases as described above.

