## Supplementary tables and figures for "Phylogeography of bar-tailed godwits: pre-LGM structure in Beringia and westward colonization of post-glacial Europe" (from another version of this preprint: version 2, 2024-05-08)

Conklin et al.

### SUPPLEMENTARY FIGURES & TABLES

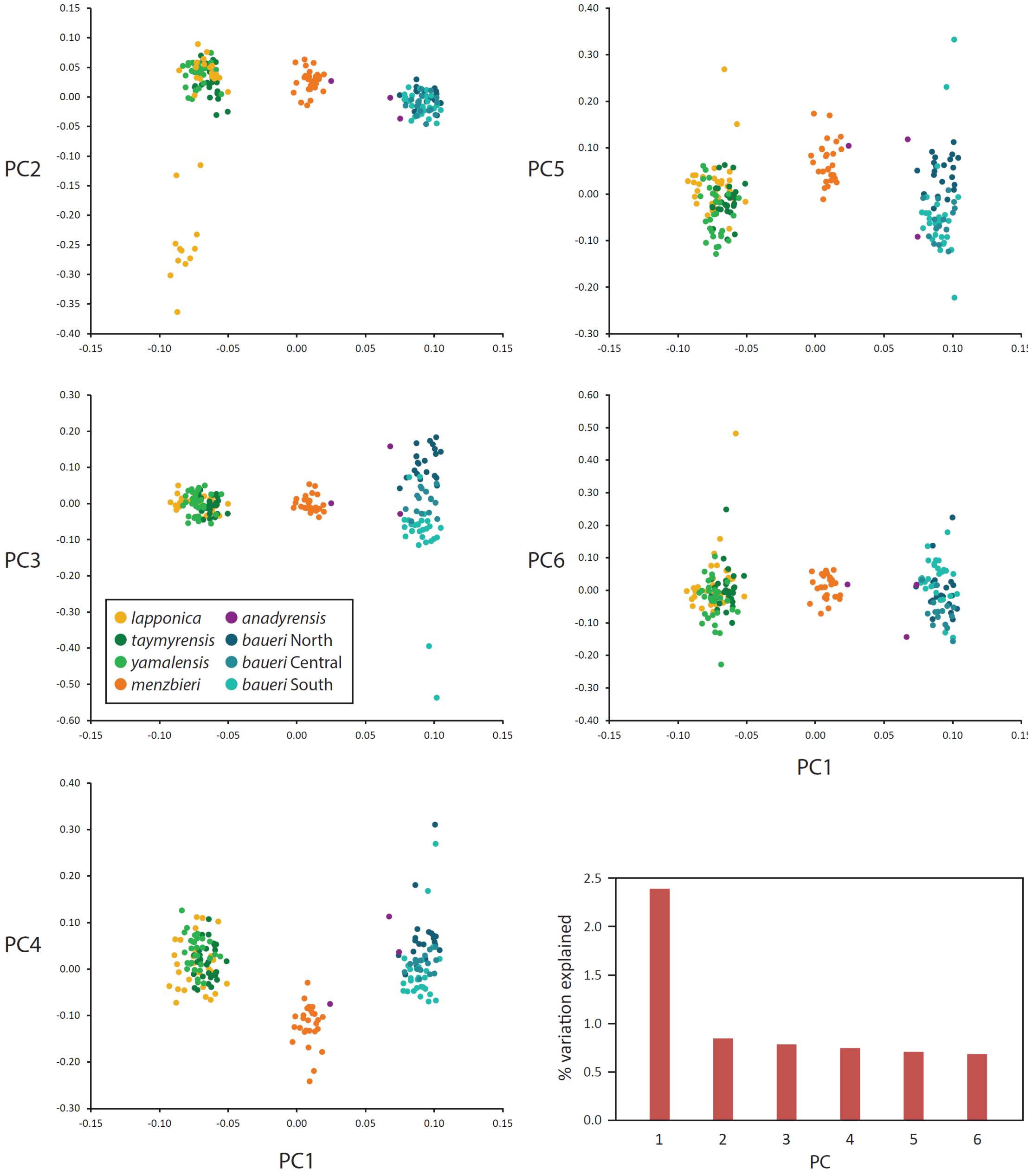


**Figure S1.** Population structure estimated by principal component analysis. The individual scores for the first principal component (PC1) are shown against the five others (PC2–6). On lower right, the scree plot indicates the proportion of genetic variance explained by each PC, with PC1–6 explaining 6.2% of total variation.

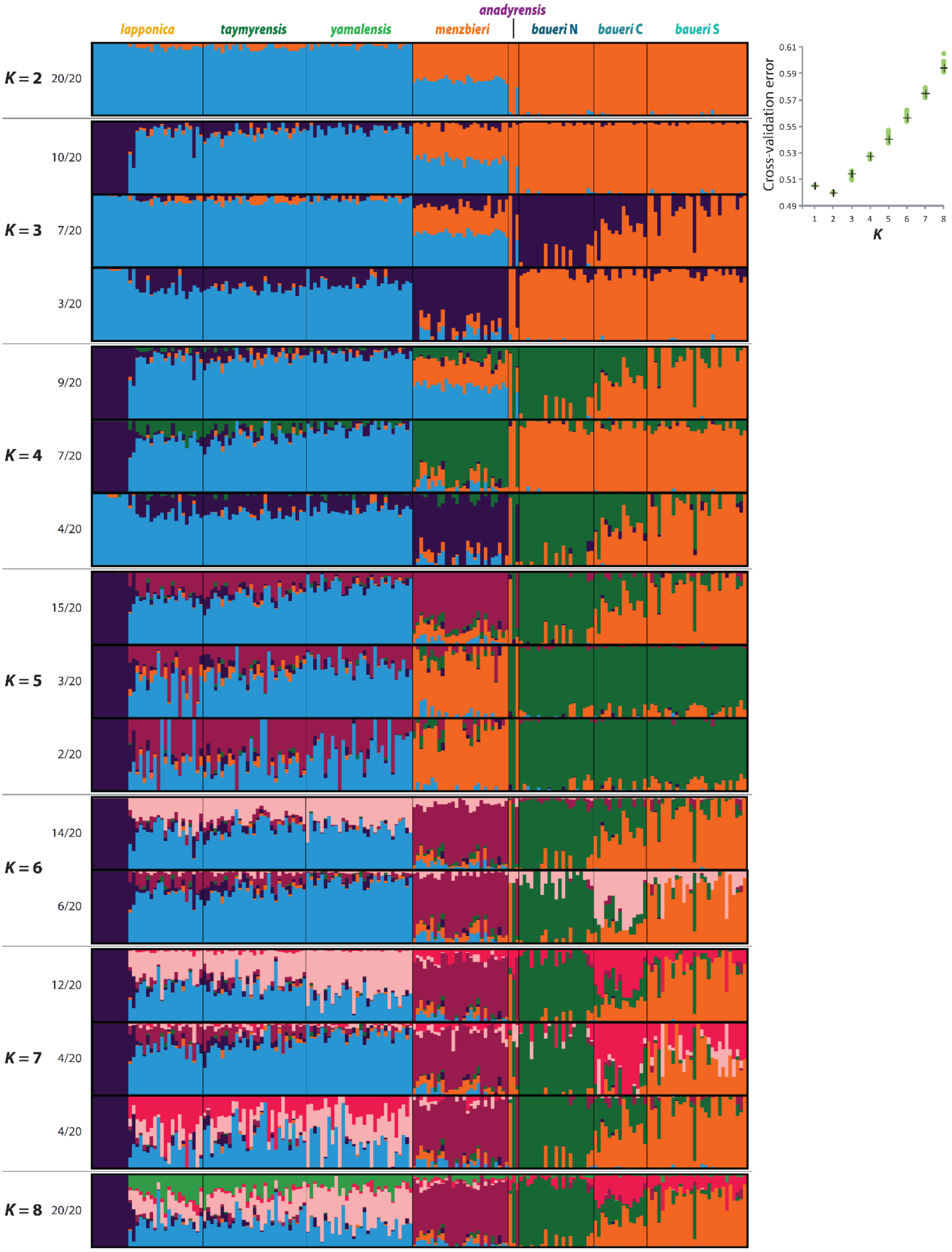


**Figure S2.** Individual genetic ancestries assigned to major and minor clusters for *K* = 2–8 estimated using ADMIXTURE. At each value of *K*, the ancestry proportions for the 184 individuals were determined by CLUMPAK summary of 20 replicate runs. Numbers on left indicate proportion of replicate runs contributing to each solution. On right, mean (black cross) and range (green dots) of CV error values for 20 replicate runs of *K* = 1–8.

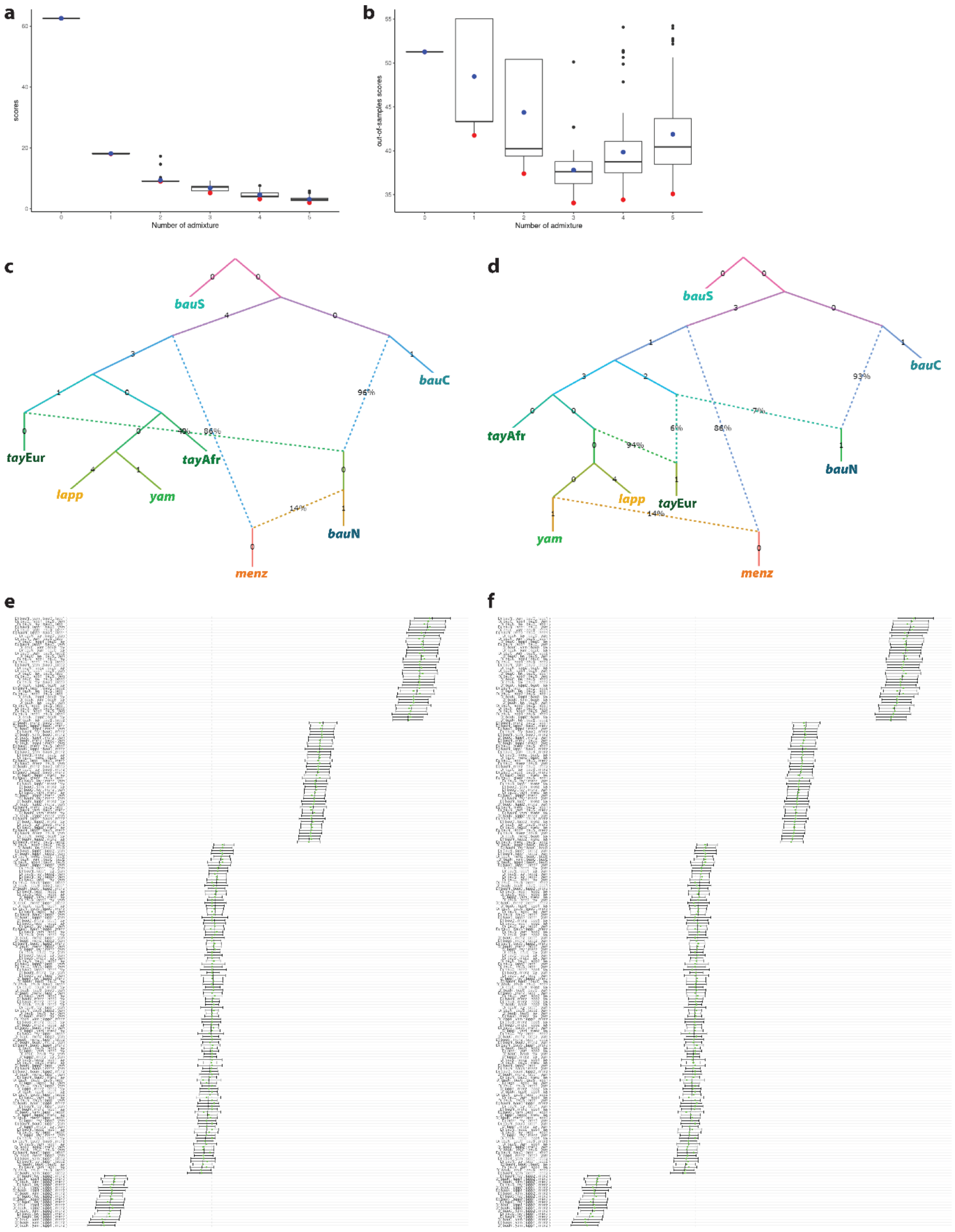


**Figure S3.** Admixture graphs estimated with ADMIXTOOLS 2.0.0: (a) Likelihood score of the admixture graphs with increasing number of admixtures (*m*). (b) Out-of-sample score values with increasing number of admixtures. (c) Best-supported admixture graph at $m = 2$. (d) Best-supported admixture graph at $m = 3$. (e,f) Plots of goodness-of-fit between observed data (green dots) and best-supported topologies at $m = 2$ and $m = 3$.

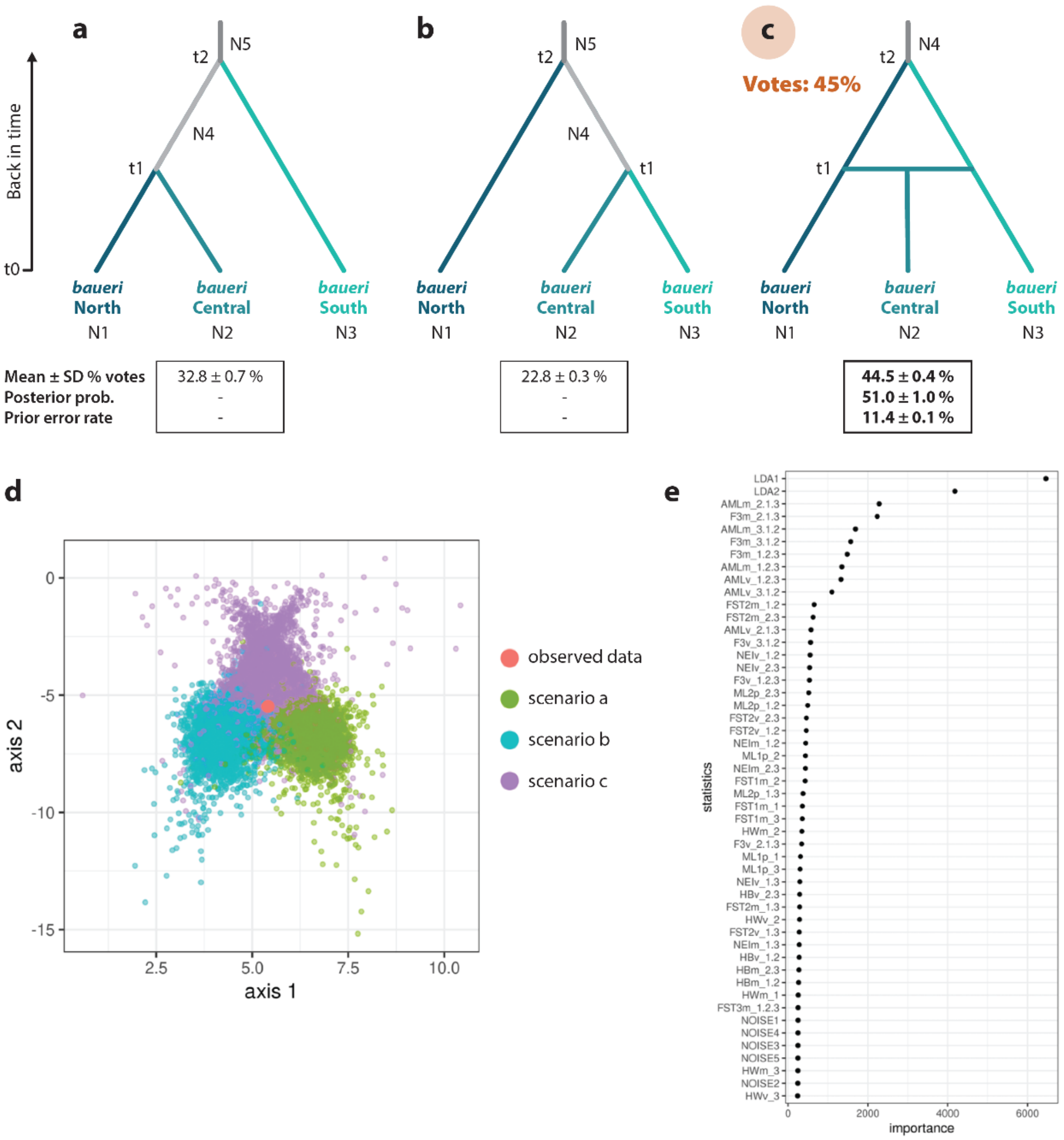


**Figure S4.** Scenarios tested in Step 1 of DIYABC-RF analysis: three possible topologies for structure within *baueri*, either simple branching (a,b) or with historical admixture (c). Extant (sampled) populations are indicated by colours; inferred historical populations are shown in grey. The best-supported scenario was (c), as inferred by the proportion of Random Forest classification votes; for this scenario, posterior probabilities, and classification error rates are also indicated. (d) Scatter plot of the first two axes of the Linear Discriminant Analysis displaying the summary statistics obtained from the simulated and observed data. (e) Contributions of the most important statistics to the Random Forest to discriminate among the competing scenarios. Acronyms on the *y*-axis are defined in Table S4. See Table S2 for further details on the model parameters.

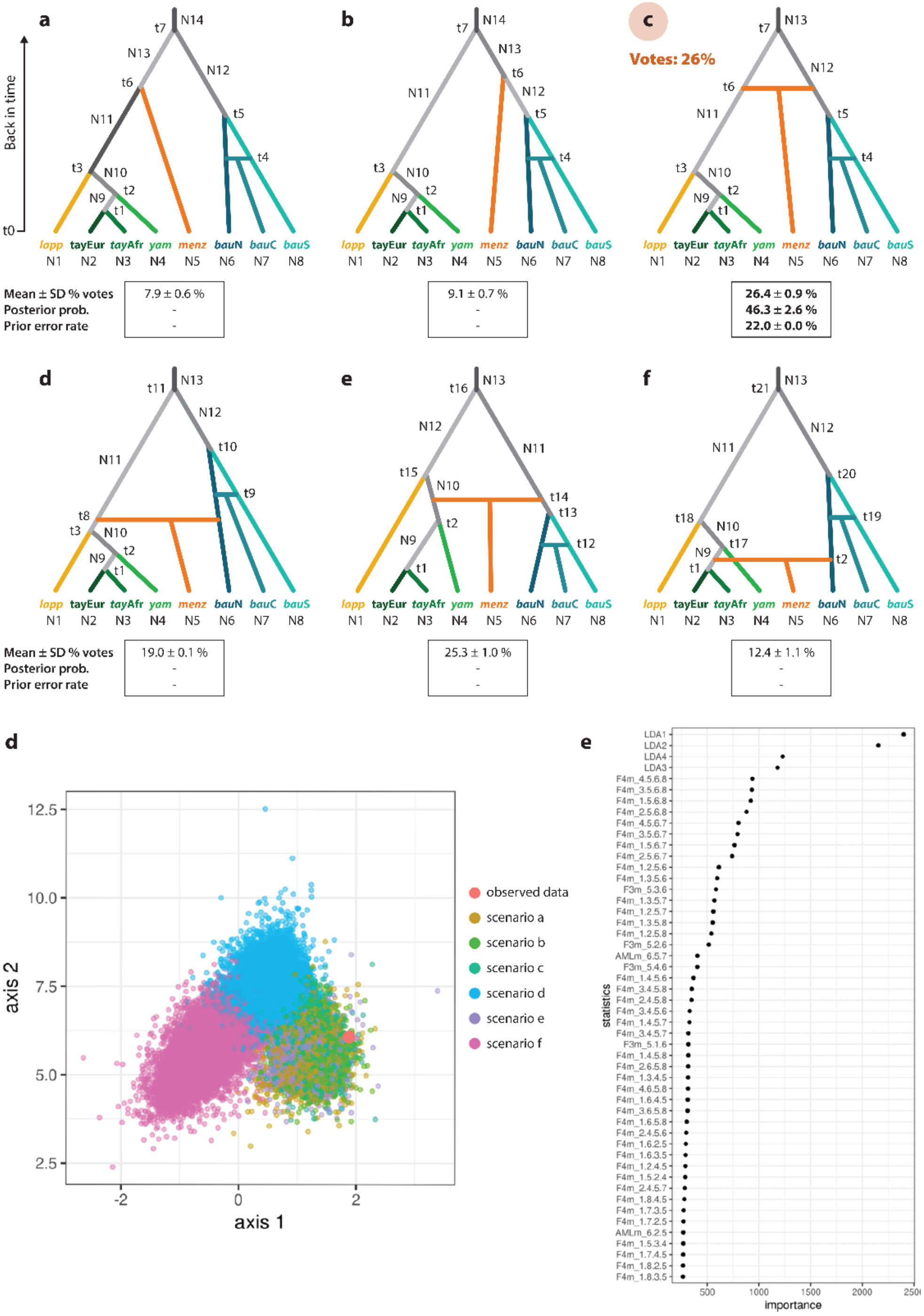


**Figure S5.** Scenarios tested in Step 2 of DIYABC-RF analysis: six possible scenarios for the origin of *menzbieri*, by either simple branching (a,b) or by admixture of the two main branches at ancient (c), intermediate (d,e) or recent (f) time periods. Extant (sampled) populations are indicated by colours; inferred historical populations are shown in grey. The best-supported scenario was (c), followed by (e), as inferred by the proportion of Random Forest classification votes; for this scenario, posterior probabilities, and classification error rates are also indicated. (g) Scatter plot of the first two axes of the Linear Discriminant Analysis displaying the summary statistics obtained from the simulated and observed data. (h) Contributions of the most important statistics to the Random Forest to discriminate among the competing scenarios. Acronyms on the *y*-axis are defined in Table S4. See Table S3 for further details on the model parameters.

**Table S1.** Sampling details for global bar-tailed godwits populations. *n* Initial = samples sequenced for *de novo* SNP discovery. *n* Final = samples included in analyses after data filtering.

| Population | Sampling location | Country | Month | Year | Type[1] | Storage[2] | Source[3] | *n* Initial | *n* Final |
|---|---|---|---|---|---|---|---|---|---|
| *lapponica* | Murmansk | Russia | Jul | 1994 | B | OE | 1 | 4 | 4 |
| | Wadden Sea | Netherlands | Aug–May | 2004–2013 | R | LE | 2 | 29 | 27 |
| *taymyrensis* | Taimyr Peninsula | Russia | Jun–Jul | 1991–2003 | B | OE | 1,3,4 | 3 | 2 |
| | Banc d'Arguin | Mauritania | Dec–Apr | 2006–2010 | R | LE | 2 | 29 | 27 |
| *yamalensis* | Barr al Hikman | Oman | Jan, Dec | 2008 | P | LE | 2 | 30 | 30 |
| *menzbieri* | Roebuck Bay, WA | Australia | Feb | 2008 | S | LQ | 5 | 11 | 11 |
| | Roebuck Bay, WA | Australia | Oct–Feb | 2013–2014 | P | LE | 6 | 19 | 16 |
| *anadyrensis* | SE Chukotka | Russia | May–Jul | 2006–2015 | B | OE | 3 | 8 | 3 |
| *baueri* North | Colville River, AK | USA | Jun–Jul | 2009–2010 | B | LQ | 5 | 18 | 18 |
| | Firth of Thames, N Island | New Zealand | Feb | 2008 | S | LQ | 5 | 1 | 1 |
| | Foxton Beach, N Island | New Zealand | Feb, Nov | 2013 | G | LQ | 7 | 3 | 2 |
| *baueri* Central | Seward Peninsula, AK | USA | Jun | 2009–2011 | B | LQ | 5 | 13 | 13 |
| | Foxton Beach, N Island | New Zealand | Feb, Nov | 2013 | G | LQ | 7 | 3 | 2 |
| *baueri* South | Yukon-Kuskokwim Delta, AK | USA | Jun | 2005–2006 | B | LQ | 5 | 15 | 15 |
| | Firth of Thames, N Island | New Zealand | Feb | 2007–2008 | S | LQ | 5 | 5 | 5 |
| | Foxton Beach, N Island | New Zealand | Feb, Nov | 2013 | G | LQ | 7 | 8 | 7 |
| | Catlins, S Island | New Zealand | Feb | 2014 | G | LQ | 7 | 1 | 1 |

| | | | | | | | | | |
|---|---|---|---|---|---|---|---|---|---|
| *baueri* unknown | Foxton Beach, N Island | New Zealand | Feb, Nov | 2013 | G | LQ | 7 | 6 | 0 |
| | Golden Bay, S Island | New Zealand | Feb | 2007 | S | LQ | 5 | 1 | 0 |
| | Catlins, S Island | New Zealand | Feb | 2014 | G | LQ | 7 | 2 | 0 |
| | | | | | | | | **209** | **184** |

[1]B Captured or collected at or near known breeding area
S Captured at non-breeding site and tracked to breeding area by satellite-telemetry
G Captured at non-breeding site and tracked to breeding area by geolocator
R Captured at non-breeding site and assigned to breeding area based on individual resight history and known population movements
P Captured at non-breeding site and assigned to breeding area based on known population movements
[2]LE Blood from live capture stored in 95% ethanol
LQ Blood from live capture stored in Queen's lysis buffer
OE Organ tissue from collected specimen stored in 95% ethanol
[3]1 Burke Museum, University of Washington, USA
2 Royal Netherlands Institute for Sea Research (NIOZ), The Netherlands
3 Zoological Museum, Moscow State University, Russia
4 Royal Ontario Museum, University of Toronto, Canada
5 Alaska Science Center, United States Geological Survey, USA
6 Groningen Institute for Evolutionary Life Sciences, University of Groningen, The Netherlands
7 Zoology & Ecology Group, Massey University, Palmerston North, New Zealand

**Table S2.** Prior distribution for each parameter of the models used in Step 1 of the DIYABC-RF analyses. N = effective population size, A = admixture rate, T = time (in generations) of population split or admixture. UN: Uniform prior distribution range. See Figure S4 for parameter specifics.

| **Parameter** | **Type** | **Prior dist. range** |
|---|---|---|
| *N1* | N | UN[10–100,000] |
| *N2* | N | UN[10–100,000] |
| *N3* | N | UN[10–100,000] |
| *N4* | N | UN[10–100,000] |
| *N5* | N | UN[10–100,000] |
| *t1* | T | UN[10–30000] |
| *t2* | T | UN[10–40000] |
| *r1* | A | UN[0.01–0.99] |
| Conditions: *t2>t1*; DRAW UNTIL | | |

**Table S3.** Prior distribution for each parameter of the models used in Step 2 of the DIYABC-RF analyses. N = effective population size, A = admixture rate, T = time (in generations) of population split or admixture. UN: Uniform prior distribution range. See Figure S5 for parameter specifics.

| **Parameter** | **Type** | **Prior dist. range** | | **Parameter** | **Type** | **Prior dist. range** |
|---|---|---|---|---|---|---|
| *t1* | T | UN[10–3000] | | *N1* | N | UN[10–100000] |
| *t2* | T | UN[100–5000] | | *N2* | N | UN[10–100000] |
| *t3* | T | UN[400–30000] | | *N3* | N | UN[10–100000] |
| *t4* | T | UN[400–30000] | | *N4* | N | UN[10–100000] |
| *t5* | T | UN[400–30000] | | *N5* | N | UN[10–100000] |
| *t6* | T | UN[400–30000] | | *N6* | N | UN[10–100000] |
| *t7* | T | UN[2000–40000] | | *N7* | N | UN[10–100000] |
| *t8* | T | UN[400–30000] | | *N8* | N | UN[10–100000] |
| *t9* | T | UN[400–30000] | | *N9* | N | UN[10–100000] |
| *t10* | T | UN[400–30000] | | *N10* | N | UN[10–100000] |
| *t11* | T | UN[2000–40000] | | *N11* | N | UN[10–100000] |
| *t12* | T | UN[400–30000] | | *N12* | N | UN[10–100000] |
| *t13* | T | UN[400–30000] | | *N13* | N | UN[10–100000] |
| *t14* | T | UN[400–30000] | | *N14* | N | UN[10–100000] |
| *t15* | T | UN[400–30000] | | *r1* | A | UN[0.01–0.99] |
| *t16* | T | UN[2000–40000] | | *r2* | A | UN[0.01–0.99] |
| *t17* | T | UN[400–30000] | | | | |
| *t18* | T | UN[400–30000] | | | | |
| *t19* | T | UN[400–30000] | | | | |
| *t20* | T | UN[400–30000] | | | | |
| *t21* | T | UN[2000–40000] | | | | |
| Conditions: *t2>t1*; *t3>t2*; *t5>t4*; *t6>t5*; *t7>t6*; *t8>t3*; *t9>t8*; *t10>t9*; *t11>t10*; *t13>t12*; *t14>t13*; *t15>t14*; *t16>t15*; *t17>t2*; *t18>t17*; *t19>t18*; *t20>t19*; *t21>t20*; DRAW UNTIL | | | | | | |

**Table S4.** Vector of summary statistics describing SNP genetic variation used in the DIYABC-RF analyses. [m|v] suffix indicates that the mean and variance (v suffix) over loci values are computed for the summary statistics.

| | |
|---|---|
| $ML_i$ | Proportion of monomorphic loci for each population ($i$), as well as for each pair, triplet and quadruplet of populations. |
| $HW_{[m\vert v]}$ and $HB_{[m\vert v]}$ | Heterozygosity within population and between pairs of populations (Hivert et al., 2018). |
| $FST_{[m\vert v]}$ | $F_{ST}$-related statistics for each population (i.e., population-specific $F_{ST}$; Weir & Goudet, 2017), for each pair, triplet, quadruplet and overall populations (when the data set includes more than four populations) (Hivert et al., 2018). |
| $f3_{[m\vert v]}$ and $f4_{[m\vert v]}$ | Patterson's allele sharing *f-statistics* for each triplet ($f_3$-statistics) and quadruplet ($f_4$-statistics) of populations (Patterson et al., 2012). |
| $Nei_{[m\vert v]}$ | Nei's (1972) distance for each pair of populations |
| $AML_{[m\vert v]}$ | Maximum likelihood coefficient of admixture computed for each triplet of populations (adapted from Choisy et al., 2004). |

**Table S5.** Descriptive statistics per population (mean ± SD) for final data set, after re-assigning 19 purported *lapponica* individuals to *taymyrensis* Europe and excluding *anadyrensis*. π = nucleotide diversity; Het = heterozygosity, $F_{IS}$ = inbreeding coefficient.

| **Population** | ***n*** | **Depth coverage** | ***π*** | **Het** | **$F_{IS}$** | **Tajima's D** |
|---|---|---|---|---|---|---|
| All | 181 | 50.06 ± 22.87 | 0.245 ± 0.145 | 0.207 ± 0.020 | 0.151 ± 0.083 | 0.765 ± 0.795 |
| *lapponica* | 12 | 57.19 ± 24.34 | 0.234 ± 0.179 | 0.198 ± 0.021 | 0.191 ± 0.085 | 0.214 ± 0.860 |
| *taymyrensis* Europe | 19 | 62.07 ± 30.93 | 0.242 ± 0.163 | 0.201 ± 0.031 | 0.180 ± 0.128 | 0.246 ± 0.835 |
| *taymyrensis* Africa | 29 | 44.39 ± 16.58 | 0.243 ± 0.157 | 0.209 ± 0.018 | 0.147 ± 0.075 | 0.329 ± 0.815 |
| *yamalensis* | 30 | 51.71 ± 18.64 | 0.242 ± 0.158 | 0.205 ± 0.016 | 0.160 ± 0.066 | 0.339 ± 0.821 |
| *menzbieri* | 27 | 43.09 ± 23.58 | 0.243 ± 0.157 | 0.215 ± 0.014 | 0.119 ± 0.059 | 0.300 ± 0.818 |
| *baueri* North | 21 | 52.32 ± 24.40 | 0.236 ± 0.167 | 0.210 ± 0.019 | 0.140 ± 0.079 | 0.277 ± 0.848 |
| *baueri* Central | 15 | 49.03 ± 21.34 | 0.235 ± 0.172 | 0.212 ± 0.017 | 0.133 ± 0.068 | 0.221 ± 0.847 |
| *baueri* South | 28 | 48.55 ± 22.81 | 0.236 ± 0.163 | 0.205 ± 0.021 | 0.160 ± 0.085 | 0.327 ± 0.846 |

**Table S6.** Complete results of $f_3$ three-population test in ADMIXTOOLS 2.0.0. The 12 scenarios indicating admixture (significant negative $f_3$ estimate) are shown in bold type.

| Scenario | Pop A | Pop B | Pop C | $f_3$ estimate | SE | $Z$ | $p$ |
|---|---|---|---|---|---|---|---|
| 1 | bauC | bauS | bauN | 0.0010 | 0.0001 | 9.78 | < 0.001 |
| 2 | bauC | bauS | lapp | 0.0013 | 0.0001 | 9.09 | < 0.001 |
| 3 | bauC | bauS | tayEur | 0.0014 | 0.0001 | 11.38 | < 0.001 |
| 4 | bauC | bauS | menz | 0.0014 | 0.0001 | 11.68 | < 0.001 |
| 5 | bauC | bauS | tayAfr | 0.0014 | 0.0001 | 10.99 | < 0.001 |
| 6 | bauC | bauS | yam | 0.0013 | 0.0001 | 10.99 | < 0.001 |
| 7 | bauC | lapp | bauN | 0.0011 | 0.0002 | 7.19 | < 0.001 |
| 8 | bauC | lapp | tayEur | 0.0083 | 0.0002 | 34.08 | < 0.001 |
| 9 | bauC | lapp | menz | 0.0048 | 0.0002 | 23.69 | < 0.001 |
| 10 | bauC | lapp | tayAfr | 0.0083 | 0.0002 | 33.38 | < 0.001 |
| 11 | bauC | lapp | yam | 0.0085 | 0.0003 | 33.81 | < 0.001 |
| 12 | bauC | tayEur | bauN | 0.0014 | 0.0001 | 10.41 | < 0.001 |
| 13 | bauC | tayEur | menz | 0.0050 | 0.0002 | 32.88 | < 0.001 |
| 14 | bauC | tayEur | tayAfr | 0.0083 | 0.0002 | 39.14 | < 0.001 |
| 15 | bauC | tayEur | yam | 0.0083 | 0.0002 | 37.42 | < 0.001 |
| 16 | bauC | menz | bauN | 0.0013 | 0.0001 | 12.37 | < 0.001 |
| 17 | bauC | menz | tayAfr | 0.0049 | 0.0002 | 30.97 | < 0.001 |
| 18 | bauC | menz | yam | 0.0049 | 0.0002 | 32.29 | < 0.001 |
| 19 | bauC | tayAfr | bauN | 0.0014 | 0.0001 | 11.02 | < 0.001 |
| 20 | bauC | tayAfr | yam | 0.0084 | 0.0002 | 38.28 | < 0.001 |
| 21 | bauC | yam | bauN | 0.0013 | 0.0001 | 9.89 | < 0.001 |
| 22 | bauN | bauC | bauS | 0.0015 | 0.0001 | 15.32 | < 0.001 |
| 23 | bauN | bauC | lapp | 0.0014 | 0.0002 | 8.60 | < 0.001 |
| 24 | bauN | bauC | tayEur | 0.0011 | 0.0001 | 8.25 | < 0.001 |
| 25 | bauN | bauC | menz | 0.0013 | 0.0001 | 11.31 | < 0.001 |
| 26 | bauN | bauC | tayAfr | 0.0012 | 0.0001 | 9.90 | < 0.001 |
| 27 | bauN | bauC | yam | 0.0012 | 0.0001 | 9.38 | < 0.001 |
| 28 | bauN | bauS | lapp | 0.0016 | 0.0001 | 13.05 | < 0.001 |
| 29 | bauN | bauS | tayEur | 0.0015 | 0.0001 | 12.50 | < 0.001 |
| 30 | bauN | bauS | menz | 0.0016 | 0.0001 | 14.09 | < 0.001 |
| 31 | bauN | bauS | tayAfr | 0.0015 | 0.0001 | 12.72 | < 0.001 |
| 32 | bauN | bauS | yam | 0.0015 | 0.0001 | 12.46 | < 0.001 |
| 33 | bauN | lapp | tayEur | 0.0082 | 0.0003 | 32.78 | < 0.001 |
| 34 | bauN | lapp | menz | 0.0050 | 0.0002 | 26.28 | < 0.001 |
| 35 | bauN | lapp | tayAfr | 0.0084 | 0.0002 | 35.71 | < 0.001 |
| 36 | bauN | lapp | yam | 0.0086 | 0.0003 | 33.84 | < 0.001 |
| 37 | bauN | tayEur | menz | 0.0048 | 0.0002 | 28.68 | < 0.001 |
| 38 | bauN | tayEur | tayAfr | 0.0081 | 0.0002 | 35.59 | < 0.001 |
| 39 | bauN | tayEur | yam | 0.0082 | 0.0002 | 35.04 | < 0.001 |
| 40 | bauN | menz | tayAfr | 0.0048 | 0.0002 | 28.71 | < 0.001 |
| 41 | bauN | menz | yam | 0.0049 | 0.0002 | 28.16 | < 0.001 |
| 42 | bauN | tayAfr | yam | 0.0082 | 0.0002 | 37.29 | < 0.001 |
| 43 | bauS | bauN | bauC | 0.0013 | 0.0001 | 12.25 | < 0.001 |
| 44 | bauS | lapp | bauC | 0.0010 | 0.0001 | 8.41 | < 0.001 |
| 45 | bauS | lapp | bauN | 0.0011 | 0.0001 | 9.69 | < 0.001 |
| 46 | bauS | lapp | tayEur | 0.0078 | 0.0002 | 36.84 | < 0.001 |
| 47 | bauS | lapp | menz | 0.0045 | 0.0002 | 26.68 | < 0.001 |
| 48 | bauS | lapp | tayAfr | 0.0080 | 0.0002 | 35.67 | < 0.001 |
| 49 | bauS | lapp | yam | 0.0082 | 0.0002 | 37.67 | < 0.001 |
| 50 | bauS | tayEur | bauC | 0.0009 | 0.0001 | 7.23 | < 0.001 |
| 51 | bauS | tayEur | bauN | 0.0012 | 0.0001 | 10.28 | < 0.001 |
| 52 | bauS | tayEur | menz | 0.0045 | 0.0001 | 34.57 | < 0.001 |
| 53 | bauS | tayEur | tayAfr | 0.0078 | 0.0002 | 40.87 | < 0.001 |
| 54 | bauS | tayEur | yam | 0.0079 | 0.0002 | 40.71 | < 0.001 |
| 55 | bauS | menz | bauC | 0.0009 | 0.0001 | 10.26 | < 0.001 |
| 56 | bauS | menz | bauN | 0.0012 | 0.0001 | 10.93 | < 0.001 |
| 57 | bauS | menz | tayAfr | 0.0045 | 0.0001 | 30.68 | < 0.001 |
| 58 | bauS | menz | yam | 0.0045 | 0.0001 | 33.21 | < 0.001 |
| 59 | bauS | tayAfr | bauC | 0.0009 | 0.0001 | 8.56 | < 0.001 |
| 60 | bauS | tayAfr | bauN | 0.0012 | 0.0001 | 10.68 | < 0.001 |
| 61 | bauS | tayAfr | yam | 0.0080 | 0.0002 | 39.22 | < 0.001 |
| 62 | bauS | yam | bauC | 0.0010 | 0.0001 | 8.45 | < 0.001 |
| 63 | bauS | yam | bauN | 0.0012 | 0.0001 | 10.57 | < 0.001 |

| | | | | | | | |
|---|---|---|---|---|---|---|---|
| 64 | lapp | bauC | bauS | 0.0116 | 0.0003 | 34.43 | < 0.001 |
| 65 | lapp | bauN | bauC | 0.0117 | 0.0003 | 34.71 | < 0.001 |
| 66 | lapp | bauN | bauS | 0.0115 | 0.0003 | 35.14 | < 0.001 |
| 67 | lapp | tayEur | bauC | 0.0046 | 0.0002 | 20.53 | < 0.001 |
| 68 | lapp | tayEur | bauN | 0.0049 | 0.0002 | 23.88 | < 0.001 |
| 69 | lapp | tayEur | bauS | 0.0047 | 0.0002 | 22.84 | < 0.001 |
| 70 | lapp | tayEur | menz | 0.0047 | 0.0002 | 25.87 | < 0.001 |
| 71 | lapp | tayEur | tayAfr | 0.0045 | 0.0002 | 27.34 | < 0.001 |
| 72 | lapp | tayEur | yam | 0.0044 | 0.0002 | 25.79 | < 0.001 |
| 73 | lapp | menz | bauC | 0.0080 | 0.0003 | 31.42 | < 0.001 |
| 74 | lapp | menz | bauN | 0.0081 | 0.0003 | 32.14 | < 0.001 |
| 75 | lapp | menz | bauS | 0.0081 | 0.0002 | 32.73 | < 0.001 |
| 76 | lapp | menz | tayAfr | 0.0046 | 0.0002 | 24.97 | < 0.001 |
| 77 | lapp | menz | yam | 0.0044 | 0.0002 | 25.16 | < 0.001 |
| 78 | lapp | tayAfr | bauC | 0.0045 | 0.0002 | 21.00 | < 0.001 |
| 79 | lapp | tayAfr | bauN | 0.0047 | 0.0002 | 23.01 | < 0.001 |
| 80 | lapp | tayAfr | bauS | 0.0046 | 0.0002 | 23.49 | < 0.001 |
| 81 | lapp | tayAfr | yam | 0.0044 | 0.0002 | 25.32 | < 0.001 |
| 82 | lapp | yam | bauC | 0.0043 | 0.0002 | 21.02 | < 0.001 |
| 83 | lapp | yam | bauN | 0.0045 | 0.0002 | 22.44 | < 0.001 |
| 84 | lapp | yam | bauS | 0.0044 | 0.0002 | 23.64 | < 0.001 |
| 85 | tayEur | bauC | bauS | 0.0076 | 0.0002 | 42.39 | < 0.001 |
| 86 | tayEur | bauC | lapp | 0.0008 | 0.0002 | 5.14 | < 0.001 |
| 87 | tayEur | bauN | bauC | 0.0077 | 0.0002 | 35.99 | < 0.001 |
| 88 | tayEur | bauN | bauS | 0.0073 | 0.0002 | 40.61 | < 0.001 |
| 89 | tayEur | bauN | lapp | 0.0006 | 0.0001 | 4.07 | < 0.001 |
| 90 | tayEur | bauS | lapp | 0.0007 | 0.0001 | 4.95 | < 0.001 |
| 91 | tayEur | menz | bauC | 0.0041 | 0.0002 | 26.46 | < 0.001 |
| 92 | tayEur | menz | bauN | 0.0040 | 0.0002 | 26.46 | < 0.001 |
| 93 | tayEur | menz | bauS | 0.0041 | 0.0001 | 29.50 | < 0.001 |
| 94 | tayEur | menz | lapp | 0.0007 | 0.0001 | 6.90 | < 0.001 |
| 95 | tayEur | menz | tayAfr | 0.0008 | 0.0001 | 8.41 | < 0.001 |
| 96 | tayEur | menz | yam | 0.0007 | 0.0001 | 8.37 | < 0.001 |
| 97 | tayEur | tayAfr | bauC | 0.0008 | 0.0001 | 6.84 | < 0.001 |
| 98 | tayEur | tayAfr | bauN | 0.0007 | 0.0001 | 8.32 | < 0.001 |
| 99 | tayEur | tayAfr | bauS | 0.0007 | 0.0001 | 7.98 | < 0.001 |
| 100 | tayEur | tayAfr | lapp | 0.0009 | 0.0001 | 9.79 | < 0.001 |
| 101 | tayEur | tayAfr | yam | 0.0008 | 0.0001 | 9.71 | < 0.001 |
| 102 | tayEur | yam | bauC | 0.0007 | 0.0001 | 6.71 | < 0.001 |
| 103 | tayEur | yam | bauN | 0.0006 | 0.0001 | 6.49 | < 0.001 |
| 104 | tayEur | yam | bauS | 0.0006 | 0.0001 | 6.62 | < 0.001 |
| 105 | tayEur | yam | lapp | 0.0010 | 0.0001 | 8.80 | < 0.001 |
| 106 | menz | bauC | bauS | 0.0031 | 0.0001 | 23.80 | < 0.001 |
| **107** | **menz** | **bauC** | **lapp** | **-0.0004** | **0.0001** | **-3.70** | **< 0.001** |
| **108** | **menz** | **bauC** | **tayEur** | **-0.0005** | **0.0001** | **-4.84** | **< 0.001** |
| 109 | menz | bauN | bauC | 0.0032 | 0.0001 | 22.88 | < 0.001 |
| 110 | menz | bauN | bauS | 0.0028 | 0.0001 | 25.87 | < 0.001 |
| **111** | **menz** | **bauN** | **lapp** | **-0.0005** | **0.0001** | **-4.26** | **< 0.001** |
| **112** | **menz** | **bauN** | **tayEur** | **-0.0004** | **0.0001** | **-4.04** | **< 0.001** |
| **113** | **menz** | **bauS** | **lapp** | **-0.0005** | **0.0001** | **-4.46** | **< 0.001** |
| **114** | **menz** | **bauS** | **tayEur** | **-0.0005** | **0.0001** | **-5.25** | **< 0.001** |
| 115 | menz | lapp | tayEur | 0.0029 | 0.0001 | 23.76 | < 0.001 |
| **116** | **menz** | **tayAfr** | **bauC** | **-0.0005** | **0.0001** | **-5.41** | **< 0.001** |
| **117** | **menz** | **tayAfr** | **bauN** | **-0.0004** | **0.0001** | **-4.30** | **< 0.001** |
| **118** | **menz** | **tayAfr** | **bauS** | **-0.0005** | **0.0001** | **-5.22** | **< 0.001** |
| 119 | menz | tayAfr | lapp | 0.0030 | 0.0001 | 21.53 | < 0.001 |
| 120 | menz | tayAfr | tayEur | 0.0028 | 0.0001 | 24.13 | < 0.001 |
| 121 | menz | tayAfr | yam | 0.0029 | 0.0001 | 23.69 | < 0.001 |
| **122** | **menz** | **yam** | **bauC** | **-0.0005** | **0.0001** | **-5.13** | **< 0.001** |
| **123** | **menz** | **yam** | **bauN** | **-0.0005** | **0.0001** | **-5.63** | **< 0.001** |
| **124** | **menz** | **yam** | **bauS** | **-0.0005** | **0.0001** | **-5.89** | **< 0.001** |
| 125 | menz | yam | lapp | 0.0032 | 0.0001 | 24.28 | < 0.001 |
| 126 | menz | yam | tayEur | 0.0029 | 0.0001 | 23.33 | < 0.001 |
| 127 | tayAfr | bauC | bauS | 0.0072 | 0.0002 | 39.04 | < 0.001 |
| 128 | tayAfr | bauC | lapp | 0.0003 | 0.0001 | 2.31 | 0.021 |
| 129 | tayAfr | bauC | tayEur | 0.0003 | 0.0001 | 3.19 | 0.0014 |
| 130 | tayAfr | bauC | menz | 0.0037 | 0.0001 | 27.59 | < 0.001 |
| 131 | tayAfr | bauN | bauC | 0.0073 | 0.0002 | 35.86 | < 0.001 |

| | | | | | | | |
|---|---|---|---|---|---|---|---|
| 132 | tayAfr | bauN | bauS | 0.0069 | 0.0002 | 38.20 | < 0.001 |
| 133 | tayAfr | bauN | lapp | 0.0001 | 0.0001 | 0.70 | 0.48 |
| 134 | tayAfr | bauN | tayEur | 0.0004 | 0.0001 | 4.19 | < 0.001 |
| 135 | tayAfr | bauN | menz | 0.0036 | 0.0001 | 26.24 | < 0.001 |
| 136 | tayAfr | bauS | lapp | 0.0002 | 0.0001 | 1.55 | 0.12 |
| 137 | tayAfr | bauS | tayEur | 0.0004 | 0.0001 | 4.13 | < 0.001 |
| 138 | tayAfr | bauS | menz | 0.0037 | 0.0001 | 29.55 | < 0.001 |
| 139 | tayAfr | lapp | tayEur | 0.0003 | 0.0001 | 2.74 | 0.0061 |
| 140 | tayAfr | lapp | menz | 0.0002 | 0.0001 | 2.20 | 0.028 |
| 141 | tayAfr | tayEur | menz | 0.0004 | 0.0001 | 4.37 | < 0.001 |
| 142 | tayAfr | yam | bauC | 0.0003 | 0.0001 | 3.16 | 0.0016 |
| 143 | tayAfr | yam | bauN | 0.0002 | 0.0001 | 2.80 | 0.0051 |
| 144 | tayAfr | yam | bauS | 0.0002 | 0.0001 | 2.55 | 0.011 |
| 145 | tayAfr | yam | lapp | 0.0004 | 0.0001 | 4.39 | < 0.001 |
| 146 | tayAfr | yam | tayEur | 0.0003 | 0.0001 | 4.52 | < 0.001 |
| 147 | tayAfr | yam | menz | 0.0002 | 0.0001 | 4.11 | < 0.001 |
| 148 | yam | bauC | bauS | 0.0079 | 0.0002 | 37.37 | < 0.001 |
| 149 | yam | bauC | lapp | 0.0007 | 0.0002 | 4.86 | < 0.001 |
| 150 | yam | bauC | tayEur | 0.0009 | 0.0001 | 8.18 | < 0.001 |
| 151 | yam | bauC | menz | 0.0043 | 0.0002 | 27.63 | < 0.001 |
| 152 | yam | bauC | tayAfr | 0.0009 | 0.0001 | 8.72 | < 0.001 |
| 153 | yam | bauN | bauC | 0.0079 | 0.0002 | 34.81 | < 0.001 |
| 154 | yam | bauN | bauS | 0.0076 | 0.0002 | 36.13 | < 0.001 |
| 155 | yam | bauN | lapp | 0.0006 | 0.0001 | 4.01 | < 0.001 |
| 156 | yam | bauN | tayEur | 0.0010 | 0.0001 | 8.81 | < 0.001 |
| 157 | yam | bauN | menz | 0.0043 | 0.0002 | 26.02 | < 0.001 |
| 158 | yam | bauN | tayAfr | 0.0009 | 0.0001 | 9.11 | < 0.001 |
| 159 | yam | bauS | lapp | 0.0007 | 0.0001 | 4.73 | < 0.001 |
| 160 | yam | bauS | tayEur | 0.0010 | 0.0001 | 9.37 | < 0.001 |
| 161 | yam | bauS | menz | 0.0044 | 0.0002 | 27.80 | < 0.001 |
| 162 | yam | bauS | tayAfr | 0.0009 | 0.0001 | 10.31 | < 0.001 |
| 163 | yam | lapp | tayEur | 0.0007 | 0.0001 | 6.47 | < 0.001 |
| 164 | yam | lapp | menz | 0.0007 | 0.0001 | 5.32 | < 0.001 |
| 165 | yam | lapp | tayAfr | 0.0007 | 0.0001 | 7.22 | < 0.001 |
| 166 | yam | tayEur | menz | 0.0009 | 0.0001 | 10.49 | < 0.001 |
| 167 | yam | tayEur | tayAfr | 0.0008 | 0.0001 | 11.36 | < 0.001 |
| 168 | yam | menz | tayAfr | 0.0009 | 0.0001 | 10.78 | < 0.001 |

81

**Table S7.** DIYABC-RF classification votes in Step 1, based on Random Forest decision trees.

| No. trees | Votes ScenA | Votes ScenB | Votes ScenC | Selected model | Posterior prob. | Prior error rate |
|---|---|---|---|---|---|---|
| 500 | 171 | 103 | 226 | C | 0.57 | 0.115 |
| 500 | 151 | 124 | 225 | C | 0.55 | 0.116 |
| 500 | 163 | 117 | 220 | C | 0.50 | 0.116 |
| 1000 | 304 | 222 | 474 | C | 0.55 | 0.115 |
| 1000 | 325 | 231 | 444 | C | 0.46 | 0.114 |
| 1000 | 324 | 243 | 433 | C | 0.52 | 0.115 |
| **2000** | **650** | **458** | **892** | **C** | **0.52** | **0.114** |
| **2000** | **645** | **458** | **897** | **C** | **0.50** | **0.114** |
| **2000** | **670** | **449** | **881** | **C** | **0.51** | **0.115** |

**Table S8.** Confusion matrix of the RF model choice analysis for Step 1 in the DIYABC-RF analysis, reporting the average ± SD classification counts for 20K simulations repeated 3 times under each model, and (right column) the average ± SD class-specific prediction error-rates.

| | **Sim. ScenA** | **Sim. ScenB** | **Sim. ScenC** | **Class error (%)** |
|---|---|---|---|---|
| **Predicted 1** | 18773.3± 4.6 | 212.3 ± 9.1 | 2259.7 ± 10.5 | 11.6 ± 0.0 |
| **Predicted 2** | 206.3 ±4.9 | 18870.3 ± 6.4 | 2271.7 ± 11.1 | 11.6 ± 0.1 |
| **Predicted 3** | 959.3 ± 0.6 | 974.3 ± 12.5 | 15572.7 ± 10.1 | 11.0 ± 0.1 |

**Table S9.** DIYABC-RF classification votes in Step 2, based on Random Forest decision trees.

| No. trees | Votes ScenA | Votes ScenB | Votes ScenC | Votes ScenD | Votes ScenE | Votes ScenF | Selected model | Posterior prob. | Prior error rate |
|---|---|---|---|---|---|---|---|---|---|
| 500 | 37 | 50 | 148 | 88 | 121 | 56 | C | 0.49 | 0.22 |
| 500 | 34 | 47 | 126 | 99 | 132 | 62 | E | 0.39 | 0.22 |
| 500 | 40 | 61 | 134 | 88 | 124 | 53 | C | 0.42 | 0.22 |
| 1000 | 72 | 93 | 248 | 197 | 249 | 141 | E | 0.43 | 0.22 |
| 1000 | 82 | 83 | 252 | 195 | 266 | 122 | E | 0.45 | 0.22 |
| 1000 | 66 | 102 | 265 | 178 | 254 | 135 | C | 0.45 | 0.22 |
| **2000** | **155** | **161** | **563** | **383** | **508** | **230** | **C** | **0.45** | **0.22** |
| **2000** | **171** | **186** | **521** | **386** | **517** | **219** | **C** | **0.45** | **0.22** |
| **2000** | **166** | **183** | **546** | **349** | **523** | **233** | **C** | **0.52** | **0.22** |
| **2000** | **157** | **188** | **526** | **369** | **488** | **272** | **C** | **0.44** | **0.22** |
| **2000** | **175** | **187** | **496** | **384** | **537** | **221** | **E** | **0.44** | **0.22** |
| **2000** | **152** | **162** | **537** | **418** | **490** | **241** | **C** | **0.47** | **0.22** |
| **2000** | **168** | **173** | **535** | **367** | **516** | **241** | **C** | **0.46** | **0.22** |
| **2000** | **137** | **185** | **519** | **387** | **493** | **279** | **C** | **0.45** | **0.22** |
| **2000** | **146** | **205** | **512** | **395** | **472** | **270** | **C** | **0.49** | **0.22** |
| **2000** | **159** | **184** | **528** | **364** | **516** | **249** | **C** | **0.44** | **0.22** |

**Table S10.** Confusion matrix of the RF model choice analysis for Step 2 in the DIYABC-RF analysis, reporting the average ± SD classification counts for 20K simulations repeated 10 times under each model, and (right column) the average ± SD class-specific prediction error-rates.

| | **Sim. ScenA** | **Sim. ScenB** | **Sim. ScenC** | **Sim. ScenD** | **Sim. ScenE** | **Sim. ScenF** | **Class error (%)** |
|---|---|---|---|---|---|---|---|
| **Predicted A** | 16048.9 ± 16.1 | 635.3 ± 8.5 | 2929.6 ± 16.1 | 295.5 ± 5.1 | 1875.4 ± 7.6 | 10.4 ± 1.0 | 26.4 ± 0.1 |
| **Predicted B** | 728.9 ± 7.1 | 16289.2 ± 18.0 | 3556 ± 13.1 | 198.9 ± 4.7 | 1915.4 ± 13.3 | 1 ± 0.0 | 28.2 ± 0.1 |
| **Predicted C** | 2354.2 ± 20.1 | 2283 ± 21.2 | 8119.8 ± 25.6 | 238.3 ± 4.3 | 1762.8 ± 13.7 | 1.5 ± 0.8 | 45.0 ± 0.1 |
| **Predicted D** | 213.7 ± 4.4 | 109.4 ± 2.1 | 331.4 ± 5.9 | 19050.3 ± 5.0 | 61.7 ± 2.7 | 212.6 ±3.7 | 4.6 ± 0.0 |
| **Predicted E** | 582.7 ± 4.0 | 690.9 ± 11.4 | 5164.5 ± 11.8 | 77.8 ± 3.1 | 14494 ± 20.7 | 7.2 ± 1.5 | 31.0 ± 0.1 |
| **Predicted F** | 0.6 ± 0.7 | 0.2 ± 0.4 | 0.7 ± 0.5 | 95.2 ± 2.9 | 0.7 ± 0.5 | 19662.3 ± 4.9 | 0.5 ± 0.0 |

**Table S11.** Posterior distribution for each parameter of the best-fitted model (scenario C) in the Step 2 analysis of the DIYABC-RF (Figures 4 and S5), based on a training reference table of 120K simulations and 500 RF decision trees. See Figure S5 and Table S3 for parameter specifics.

| Parameter | Mode | Mean | Median | Quantile 0.05 | Quantile 0.95 |
|---|---|---|---|---|---|
| *N1* | | 67193.5 | 67542.3 | 34255.0 | 97193.0 |
| *N2* | | 35181.6 | 30392.3 | 10090.0 | 80575.6 |
| *N3* | | 53501.4 | 53171.7 | 10812.0 | 97071.3 |
| *N4* | | 33835.1 | 21755.6 | 8178.2 | 87650.2 |
| *N5* | | 59716.8 | 59658.4 | 23477.6 | 96888.0 |
| *N6* | | 36095.9 | 28673.8 | 5127.6 | 91720.9 |
| *N7* | | 52116.0 | 51842.0 | 18568.0 | 91798.0 |
| *N8* | | 17145.3 | 11903.0 | 4129.8 | 59378.6 |
| *r1* | | 0.58 | 0.60 | 0.09 | 0.95 |
| *r2* | | 0.52 | 0.52 | 0.13 | 0.91 |
| *T1* | 564 | 802.5 | 715.0 | 165.0 | 1701.0 |
| *T2* | 656 | 718.2 | 675.0 | 239.6 | 1313.0 |
| *T3* | 2053 | 2221.1 | 2054.2 | 1136.0 | 3613.6 |
| *T4* | 1452 | 2048.7 | 1787.0 | 561.0 | 4245.0 |
| *T5* | 1540 | 1746.2 | 1583.4 | 867.3 | 3066.5 |
| *T6* | 2749 | 3723.8 | 3447.7 | 1672.9 | 6565.0 |
| *T7* | 4776 | 6593.6 | 6296.0 | 2746.5 | 11639.6 |
| **Time converted from generations to years assuming 8 yr per generation** | | | | | |
| *T1yr* | 4508 | 6420.2 | 5720.0 | 1320.0 | 13608.0 |
| *T2yr* | 5248 | 5745.7 | 5400.0 | 1917.0 | 10504.0 |
| *T3yr* | 16424 | 17768.6 | 16433.6 | 9088.0 | 28908.8 |
| *T4yr* | 11613 | 16389.4 | 14296.0 | 4488.0 | 33960.0 |
| *T5yr* | 12317 | 13969.3 | 12666.8 | 6938.2 | 24532.0 |
| *T6yr* | 21988 | 29790.2 | 27581.9 | 13383.0 | 52520.0 |
| *T7yr* | 38211 | 52749.0 | 50367.6 | 21971.8 | 93116.8 |
